# An emergent disease-associated motor neuron state precedes cell death in a mouse model of ALS

**DOI:** 10.1101/2025.08.21.671404

**Authors:** Olivia Gautier, Jacob A. Blum, Thao P. Nguyen, Sandy Klemm, Mai Yamakawa, Nasa Sinnott-Armstrong, Yi Zeng, Chung-ha O. Davis, Juliane Bombosch, Lisa Nakayama, Kevin A. Guttenplan, Derek Chen, Arwa Kathira, Luke Zhao, Jessica E. Rexach, William J. Greenleaf, Aaron D. Gitler

**Author notes:** These authors contributed equally (Olivia Gautier, Jacob A. Blum).

## Abstract

To uncover molecular determinants of motor neuron degeneration and selective vulnerability in amyotrophic lateral sclerosis (ALS), we generated longitudinal single-nucleus transcriptomes and chromatin accessibility profiles of spinal motor neurons from the SOD1-G93A ALS mouse model. Vulnerable alpha motor neurons showed thousands of molecular changes, marking a transition into a novel cell state we named ‘disease-associated motor neurons’ (DAMNs). We identified transcription factor regulatory networks that govern how healthy cells transition into DAMNs as well as those linked to vulnerable and resistant motor neuron subtypes. Using spatial transcriptomics, we found reactive glia located near motor neurons early in disease, suggesting early signaling events between motor neurons and glia. Finally, we found that the human orthologs of genomic regions with differential accessibility in SOD1-G93A alpha motor neurons are enriched for single nucleotide polymorphisms associated with human ALS, providing evidence that the genetic underpinnings of motor neuron vulnerability are conserved.

## Introduction

Amyotrophic lateral sclerosis (ALS) is a progressive disorder distinguished by degeneration of motor neurons in the brain and spinal cord. Motor neuron loss impairs mobility, dexterity, speech and swallowing, and ultimately causes fatal paralysis. ALS is a cruelly rapid disease; most patients survive for just three to five years after diagnosis.^1^ Effective therapies depend on defining the cellular and molecular drivers of motor neuron degeneration. Early insight into the etiology of motor neuron loss has come from the discovery of mutations that cause inherited forms of ALS. The first ALS gene identified was *SOD1*, which encodes the antioxidant enzyme superoxide dismutase.^2^ *SOD1* mutations cause ∼20% of inherited ALS cases, constituting ∼2% of total cases.^2–4^ In these cases and in mouse models of the disease, mutant SOD1 misfolds, aggregates, and causes disease via a toxic gain-of-function that is incompletely understood.^5–11^

Expression of the ALS-associated variant *SOD1-G93A* in mice recapitulates key features observed in ALS patients, providing a useful model of ALS.^7^ In both SOD1-G93A mice and ALS patients, not all spinal motor neurons are equally susceptible to degeneration—alpha motor neurons are vulnerable, while gamma and visceral motor neurons are relatively resilient.^12–14^ Even among alpha motor neurons, the larger, fast-firing ones that innervate fast-twitch muscle fibers are most vulnerable to degeneration.^15–19^ Interestingly, this selective motor neuron vulnerability occurs despite ubiquitous *SOD1* expression.^7,20,21^

Although spinal motor neuron subtypes exhibit differing susceptibilities to degeneration in disease, most prior studies analyze motor neurons in bulk, treating motor neurons as a single population that represents the ensemble average of motor neuron subtypes present at a particular time point.^22–25^ Thus, to provide insight into motor neuron subtype-selective vulnerability, we performed longitudinal single-nucleus RNA and Assay for Transposase-Accessible Chromatin (ATAC) sequencing, along with spatial transcriptomics, to generate comprehensive transcriptomic and epigenomic profiles of vulnerable and resistant spinal motor neurons in the SOD1-G93A mouse. We discovered a population of alpha motor neurons that exhibited an extensive disease-associated gene expression and chromatin accessibility signature, which we have named ‘disease-associated motor neurons’ or DAMNs. We used the ATAC and RNA data to nominate transcription factor drivers of the DAMN state transition and mediators of motor neuron subtype-selective vulnerability. At early stages of disease, we identified reactive glial cells adjacent to motor neurons, suggesting that motor neurons may contribute to early glial cell activation. Finally, we provide evidence that the human orthologs of regions with differentially accessibility in SOD1-G93A alpha motor neurons are enriched for single nucleotide polymorphisms (SNPs) associated with human ALS, providing insights into the genetic basis of ALS and its connection to sporadic forms of the disease in humans.

## Results

### Single-nucleus RNA sequencing and spatial transcriptomics of the SOD1-G93A mouse spinal cord

To investigate the mechanisms by which motor neurons degenerate and provide insight into motor neuron selective vulnerability in ALS, we performed single nucleus RNA sequencing (snRNA-seq) of the SOD1-G93A mouse spinal cord. Motor neurons are rare in the spinal cord ^26,27^, so we used an enrichment strategy to obtain nuclei from cholinergic cells, which include all spinal motor neurons and some interneuron subtypes^28^ (Figure 1A). We then pooled nuclei from cholinergic cells and non-cholinergic cells to define the transcriptional changes that occur in other important, non-cholinergic cells of the spinal cord (e.g., interneurons, astrocytes, microglia, etc.) that may play non-cell autonomous roles in motor neuron degeneration (Figure 1A). Defects in skeletal motor neuron physiology emerge even prior to overt phenotypic onset^15,16,29^, so we collected timepoints from early-stage (∼P65), mid-stage (∼P100), and end-stage (∼P125) animals, which exhibit progressive hind limb paralysis (Figure 1A). We also profiled cells from age-matched, non-transgenic control mice (Figure 1A). In total, we transcriptionally profiled >115,000 high-quality nuclei from these four conditions (Figure S1A, Figure 1B).

**Figure 1:**
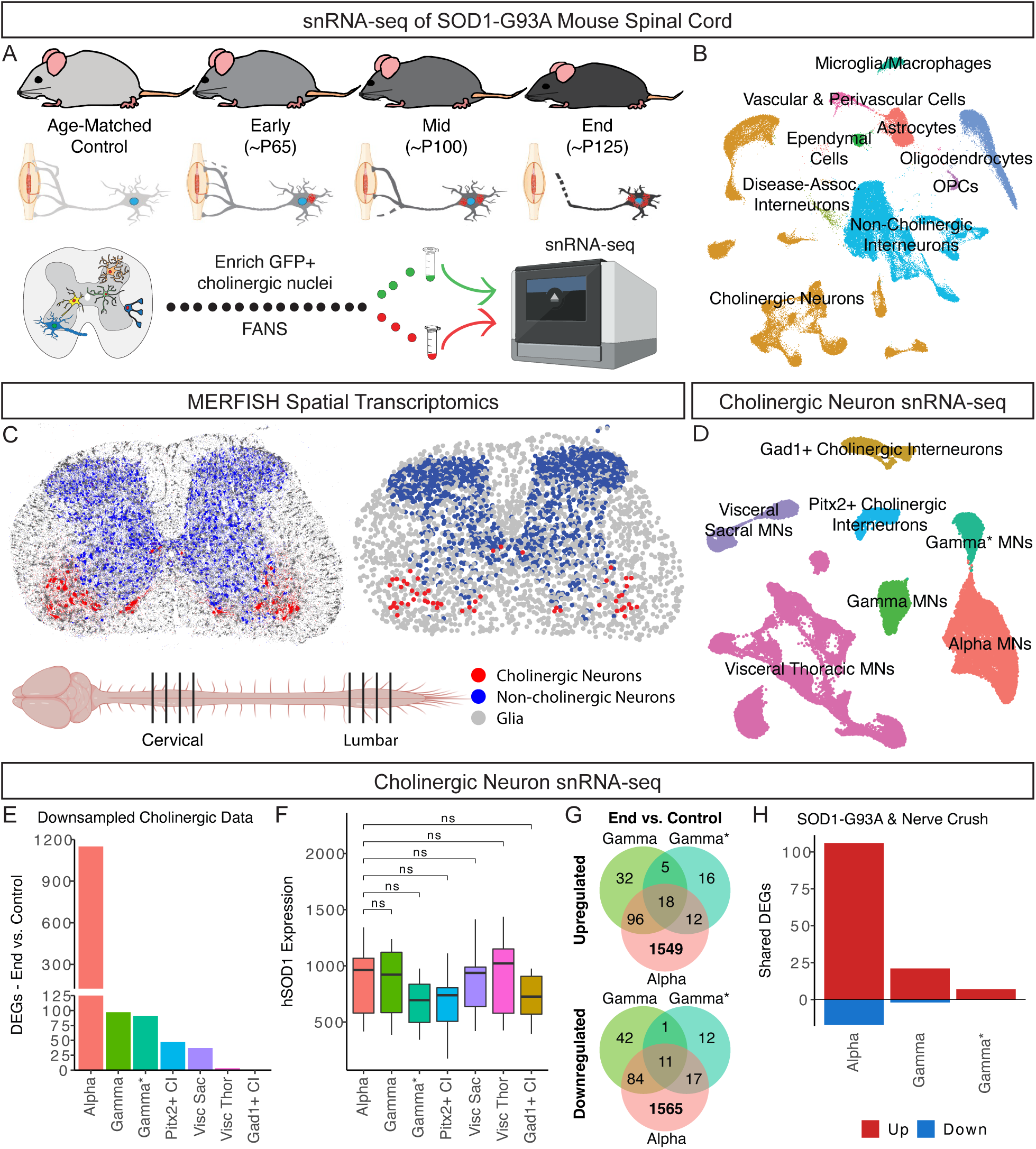
Single-cell transcriptional profiling of the SOD1-G93A mouse spinal cord. **(A)** Spinal cords were collected from early, mid, and end-stage SOD1-G93A mice and age-matched controls and processed for snRNA-seq. Cholinergic nuclei were GFP+ and enriched using fluorescence-activated nuclear sorting (FANS). **(B)** UMAP representation of all snRNA-seq nuclei labeled by major cell class. **(C)** Cervical and lumbar spinal cord sections from SOD1-G93A and control mice were processed for MERFISH spatial transcriptomics. Representative data from a control lumbar section are shown: (left) each dot represents a transcript, with markers for cholinergic neurons (*Chat*, *Prph*, *Slc5a7*, red), non-cholinergic interneurons (*Gad1*, *Slc17a6*, *Slc6a5*, blue), and glia (*Aldh1l1*, *Aqp4*, *Cx3cr1*, *Trem2*, *Mog*, gray); (right) each dot represents an individual cell colored by the same categories (cholinergic neurons, non-cholinergic interneurons, and glia). Other cell types are not shown. **(D)** UMAP representation of cholinergic neurons labeled by cholinergic neuron type. **(E)** Bar plot showing the number of differentially expressed genes between end-stage SOD1-G93A and control nuclei for each cholinergic neuron type using downsampled data (Wald test with Benjamini-Hochberg correction; padj < 0.01). **(F)** Box plots showing the average expression of the hSOD1-G93A transgene (counts per million, CPM) across cholinergic neuron subtypes. Each point represents the average expression from a single experiment (n = 11-15 experiments with >10 cells per group). Pairwise comparisons were performed between alpha motor neurons and each other subtype using Welch’s t-test with Benjamini-Hochberg false discovery rate correction. Asterisks denote statistical significance: n.s. = not significant. **(G)** Venn diagrams of genes significantly upregulated (top) or downregulated (bottom) in alpha, gamma, and gamma* motor neurons at end-stage compared to control (Wald test with Benjamini-Hochberg correction; padj < 0.01). **(H)** Overlap of differentially expressed genes from SOD1-G93A motor neurons (from G) and spinal cholinergic neurons after sciatic nerve crush from Shadrach et al., 2021.

We clustered nuclei and annotated them using expression of known marker genes^28,30^ (Figure S1A-S1B, Figure 1B), resulting in ∼39% of nuclei coming from cholinergic neurons, including motor neurons, and the rest coming from other spinal cord cell types. To validate our sequencing results and provide spatial context to our data, we performed multiplexed error-robust fluorescence *in situ* hybridization (MERFISH)^31^ to simultaneously probe expression of 140 genes in control and SOD1-G93A mouse spinal cords throughout disease progression (Figure 1C, Figure S1C). Clustering revealed expected major cell classes and spatial location of cholinergic neurons (Figure S1C-S1D, Figure 1C).

### Gene expression changes in spinal motor neurons during neurodegeneration

To identify gene expression changes with disease in vulnerable and resistant motor neurons, we first subclustered the cholinergic nuclei from the snRNA-seq and MERFISH data and annotated the resulting clusters using known marker genes^28,32^ (Figure S2A-S2E, Figure 1D). These clusters correspond to various subtypes of skeletal motor neurons, visceral motor neurons, and cholinergic interneurons (Figure 1D, Figure S2E). We performed pseudobulk differential expression analysis with the snRNA-seq data to find differentially expressed genes (DEGs) between control and end-stage disease conditions for all cholinergic neuron types (Figure 1E). Because the different types of cholinergic neurons are not equally abundant in the spinal cord or in the sequencing data, we downsampled the data to control for differences in the number of nuclei for each cholinergic type (see Methods). Among cholinergic neurons, alpha motor neurons had the greatest number of DEGs by far, over an order of magnitude more than any of the other cholinergic neuron types, which all showed a relatively minor transcriptional response to hSOD1-G93A expression (Figure 1E). The hSOD1-G93A transgene was expressed at equivalent levels in alpha motor neurons and the other cholinergic neuron types (Figure 1F), suggesting that the large transcriptional response to disease in alpha motor neurons was not due to disproportionate levels of mutant SOD1 RNA.

Next, we compared DEGs with disease among vulnerable and resistant skeletal motor neurons to identify common and divergent gene expression changes (Figure 1I, Table S1A-S1C). We first focused on DEGs shared among all skeletal motor neuron subtypes with disease (18 upregulated and 11 downregulated) (Figure 1G, Table S1A-S1C). Of the shared upregulated genes, we found the regeneration-associated transcription factors (TFs) *Sox11* and *Stat3*, which have both been shown to play a role in peripheral axon regeneration *in vivo*^33–36^, as well as the known injury-induced genes *Socs3* and *Abca1*^24,36,37^ (Figure 1G, Table S1A-S1C).

These findings prompted us to further assess which gene expression changes in the SOD1-G93A mouse may be part of a pro-regeneration transcriptional response. To do so, we compared our results to bulk translatome data from regenerating motor neurons after sciatic nerve crush.^24^ Of the upregulated disease genes in resistant motor neurons, ∼14% are also upregulated during regeneration after nerve injury (21/151 in gamma and 7/51 in gamma*), and in alpha motor neurons, ∼6% of upregulated genes are shared (106/1,675) (Figure 1H, Table S1D). Some of these genes (e.g., *Gal*, *Adcyap1*, *Gap43*/*Basp1*, *Sprr1a*, *Atf3*) play a role in the regenerative process of peripheral neurons.^38–42^ In contrast, there was little overlap between downregulated transcripts after nerve injury and those that change in disease for alpha motor neurons (17/1,677) and gamma motor neurons (2/138) (Figure 1H, Table S1D). These results suggest that all skeletal motor neurons undergo transcriptional changes associated with regeneration in the SOD1-G93A mouse, but these changes are ultimately insufficient to prevent degeneration and death of alpha motor neurons.

### A disease-associated motor neuron (DAMN) signature in alpha motor neurons

To identify changes in alpha motor neuron subtypes with disease, we subclustered the alpha motor neuron transcriptomes into 12 distinct groups (Figure 2A). Strikingly, we observed the emergence of a new transcriptionally distinct alpha motor neuron cluster with disease (cluster 2; Figure 2B-2C), which was present at ∼4% at early-stage and increased in representation to ∼35% at the end-stage of disease (Figure 2D). We have named cells in this new cluster ‘disease-associated motor neurons’ or DAMNs. Hierarchical clustering separated DAMNs from all other SOD1-G93A alphas, which grouped with controls, indicating that many non-DAMNs remain control-like even at end-stage (Figure 2E). DAMNs have thousands of DEGs when compared to non-DAMNs (Figure 2F, Table S1E), which perforce warrants classification as a new cell state. While the DAMN cluster increases in relative abundance with disease (Figure 2D), clusters 7 and 9 are significantly depleted (Figure S3A). Cluster 7 is marked by *Sema3e* and *Cdh8*, and may correspond to gluteus and shoulder-innervating alpha motor neurons^28,43,44^. Cluster 9 is marked by genes associated with fast-firing (FF) alpha motor neurons (Figure S3C)^28,45^ and lacks expression of slow-firing (SF) marker genes (Figure S3D)^28,46^, and its reduction in disease is consistent with the known preferential vulnerability of FF alpha motor neurons^15–18^.

**Figure 2:**
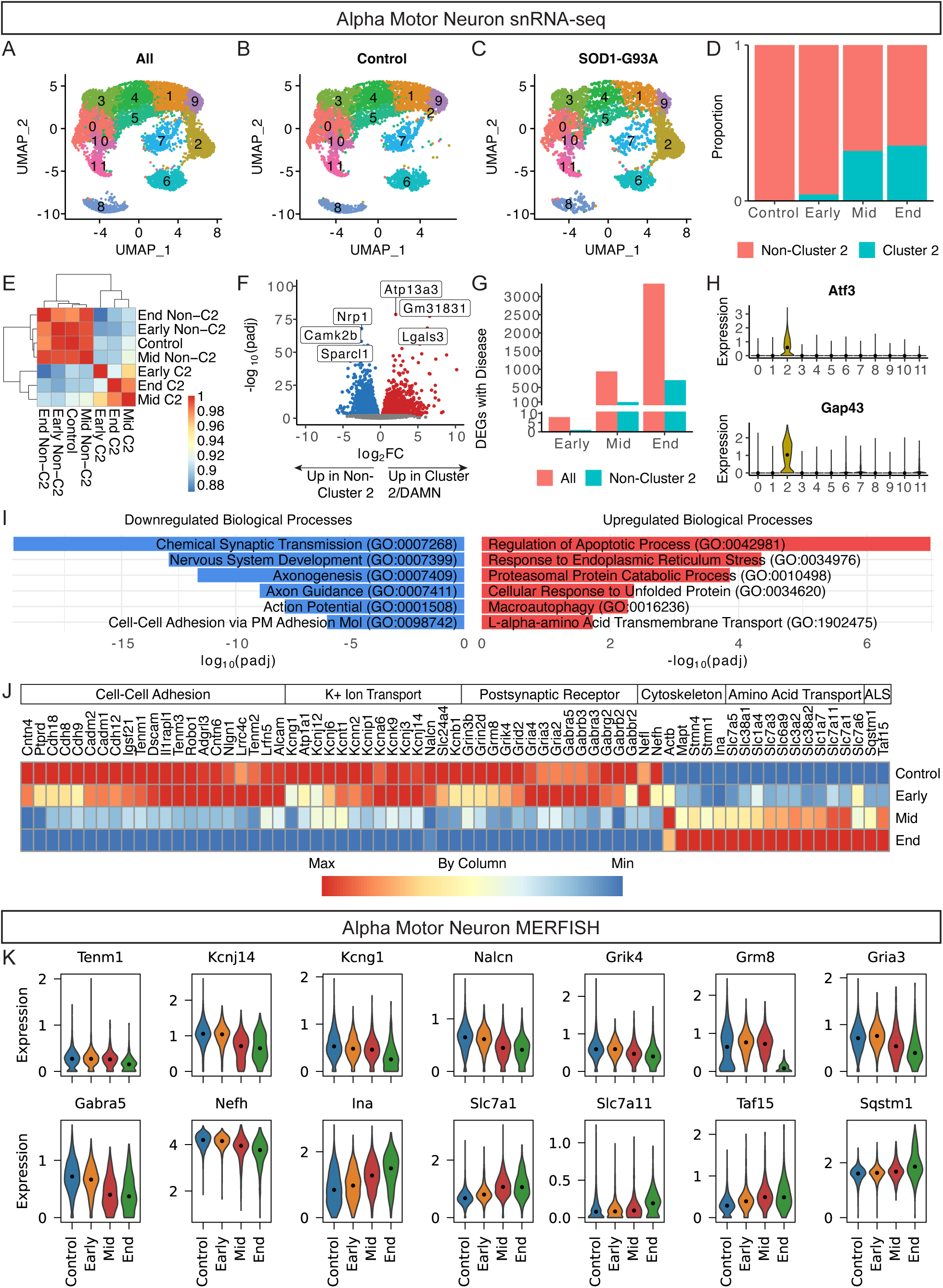
Transcriptional signature of disease-associated alpha motor neurons (A-C) UMAP representations of all **(A)**, control **(B)**, and disease **(C)** alpha motor neurons labeled by cluster. **(D)** Stacked bar plots showing the proportion of alpha motor neurons in disease-associated cluster 2 or other clusters in control and disease conditions. **(E)** Heatmap of Pearson correlations between average expression profiles of alpha motor neuron groups. Groups were defined by disease stage (Control, Early, Mid, End) and cluster identity (C2 or non-C2). Correlations were computed using the top 2,000 variable genes, and hierarchical clustering was applied to rows and columns. **(F)** Volcano plot showing genes significantly upregulated (red) or downregulated (blue) in cluster 2/DAMNs compared to cells in other clusters (Wald test with Benjamini-Hochberg correction; padj < 0.01). **(G)** Bar plots of the number of differentially expressed genes in alpha motor neurons between early, mid, or end-stage and control conditions with or without the inclusion of disease-associated cluster 2 (Wald test with Benjamini-Hochberg correction; padj < 0.01). **(H)** Violin plots showing the log-normalized expression of *Atf3* (top) and *Gap43* (bottom) in alpha motor neuron clusters. Dots mark the median values. **(I)** Bar plots showing log_10_(padj) or −log_10_(padj) for selected downregulated (left) and upregulated (right) GO biological processes in DAMNs. **(J)** Heatmap showing average expression levels of selected differentially expressed genes throughout disease progression in alpha motor neurons. Data are min/max-normalized by column. **(K)** Violin plots showing the log-normalized expression of select differentially expressed genes in alpha motor neurons in control and disease conditions by MERFISH (two-sided Mann-Whitney U test between control and end-stage with Benjamini-Hochberg correction; FDR < 0.01). Dots mark the median values.

As the proportion of DAMNs increased with disease progression, the number of DEGs between disease and control cells also increased (Figure 2G). We also detected disease-associated DEGs when we excluded the DAMN cluster from the analysis (Figure 2G), implying that some early transcriptional changes occur even before alpha motor neurons enter the DAMN cluster. To identify these pre-DAMN changes, we performed differential expression analysis between the control and disease conditions separately for FF alpha motor neurons (clusters 9 and 1) and SF alpha motor neurons (clusters 0 and 10) (Figure S3C). We observed a greater number of gene expression changes in FF alpha motor neurons with disease (213 downregulated and 237 upregulated) than in SF ones (23 downregulated and 37 upregulated), with an overlap of 11 downregulated and 13 upregulated genes between the two groups (Figure S3E, Table S1F-S1G). The overlapping upregulated genes include *Sox11*, *Etv4*, and *Atf3* (Figure S3E-S3F), among other genes, suggesting that the pre-DAMN transcriptional response is at least partially protective. In disease, FF alpha motor neurons exhibit downregulation of genes involved in potassium ion transport and extracellular matrix organization, and upregulation of genes involved in amino acid transport, negative regulation of apoptosis, and integrated stress response signaling (Figure S3G-S3H, Table S1H-S1I), among other processes, prior to entrance into the DAMN state.

The transcriptional changes in DAMNs compared to non-DAMNs include higher expression of known markers of neuronal denervation and axon regrowth, *Atf3* and *Gap43*, and genes involved in apoptosis (Figure 2H-I, Table S1E, Table S1J), suggesting that DAMNs correspond to denervating, degenerating, and ultimately dying alpha motor neurons. DAMNs are also characterized by an upregulation of genes involved in response to ER stress/unfolded protein, proteasomal protein catabolism, autophagy, and amino acid transport, among other processes, as well as an increase in ALS-associated genes (e.g., *Sqstm1*, *Taf15*) (Figure 2I-2J, Table S1E, Table S1J). Additionally, DAMNs exhibit a marked downregulation of genes involved in synaptic transmission (including both pre- and post-synaptic genes), axonogenesis/axon guidance, potassium ion transport/action potential, and cell-cell adhesion, among other processes (Figure 2I-J, Table S1E, Table S1K). We validated a subset of these disease-associated transcriptional changes with MERFISH (Figure 2K, Figure S3I, Table S1L-S1M).

Together, these data indicate that alpha motor neurons do not enter the DAMN state all at once; even at later stages of disease, there are SOD1-G93A alpha motor neurons that remain transcriptionally closer to control alpha motor neurons than to DAMNs (Figure 2A-2E). We also identified early, pre-DAMN transcriptional changes, especially in FF alpha motor neurons, and widespread changes in the DAMN state (Figure SE, Figure 2F). These findings mirror human ALS, where relatively unaffected motor pools can coexist with markedly affected ones, and provide a gene-expression-based view of disease onset at the cellular level. To uncover potential strategies for preserving alpha motor neurons in ALS, we next sought to define the upstream epigenomic mechanisms that may contribute to their transition into the DAMN state.

### Paired single-nucleus ATAC and RNA sequencing of spinal motor neurons during neurodegeneration

To capture epigenomic changes during ALS, we profiled chromatin accessibility and gene expression using paired snATAC-seq and snRNA-seq (multiome sequencing) from the mouse SOD1-G93A spinal cord (Figure 3A). We applied a similar strategy as above to sequence substantial numbers of motor neurons and other cell types, yielding ∼27,000 high-quality multiomic profiles from control mice as well as early and mid/end-stage SOD1-G93A mice (Figure 3A-3B). Of these, ∼54% came from cholinergic neurons, including motor neurons, based on clustering and cell type annotations (Figure S4A-S4B, Figure 3B). This dataset served as the foundation for characterizing epigenomic changes with disease and across cell types.

**Figure 3:**
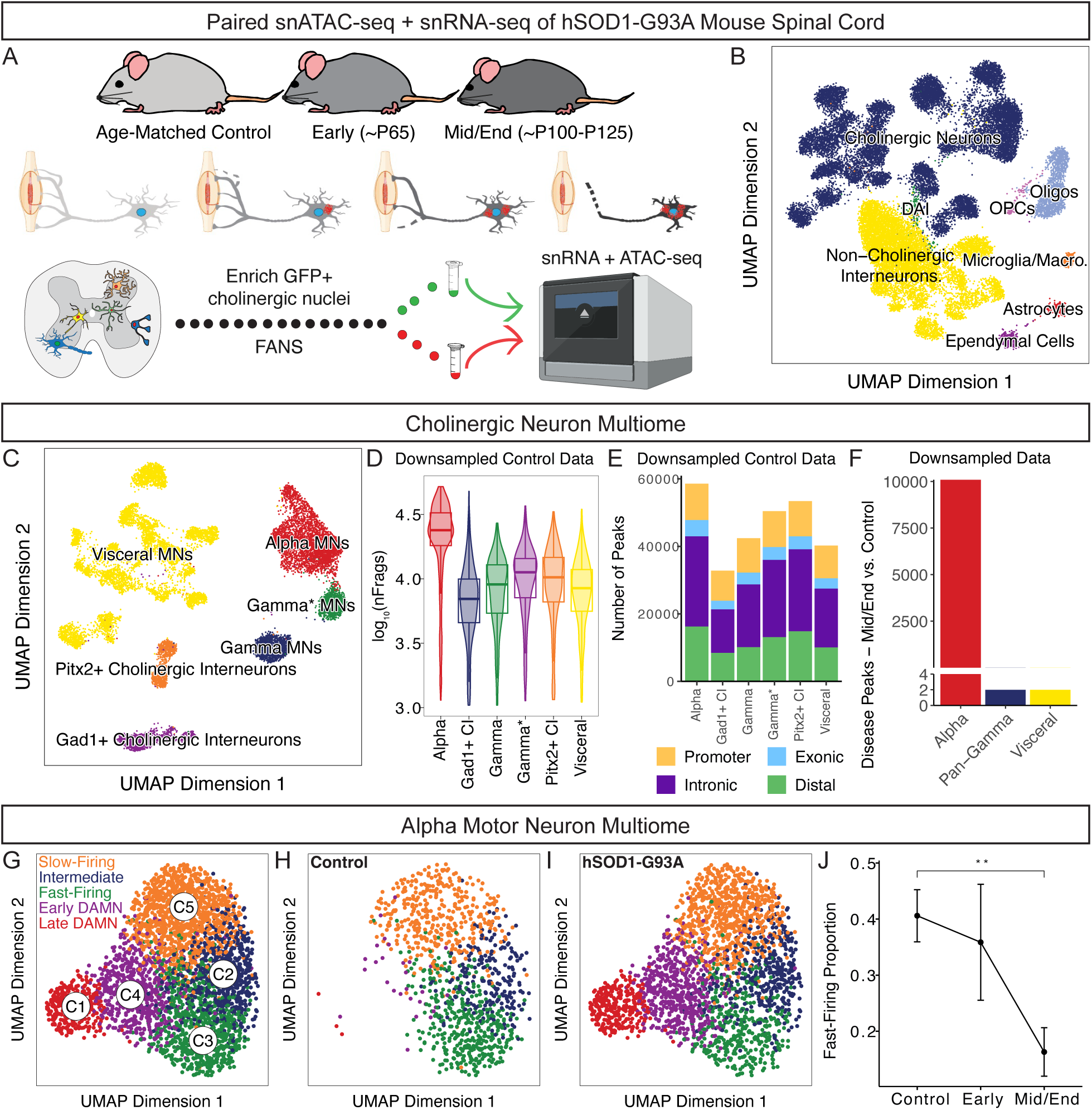
Chromatin accessibility landscapes of spinal cord cell types in the SOD1-G93A mouse. **(A)** Spinal cords were collected from early and mid/end-stage SOD1-G93A mice as well as age-matched control mice and processed for paired snATAC/snRNA-seq. Nuclei from cholinergic neurons were GFP+ and enriched using FANs. **(B)** UMAP representation of all paired snATAC/snRNA-seq nuclei labeled by major cell class. **(C)** UMAP representation of cholinergic neurons labeled by cholinergic neuron type. **(D)** Violin plots showing the distribution of ATAC-seq fragment counts (nFrags) across cholinergic neuron types in downsampled control data with box plots overlaid. **(E)** Stacked bar plots showing the number and distribution of ATAC-seq peaks across genomic annotations (Promoter, Exonic, Intronic, Distal) for each cholinergic neuron type in downsampled control data. **(F)** Bar plot showing the number of differentially accessible peaks between mid/end-stage and control nuclei for alpha, pan-gamma, and visceral motor neurons using downsampled data (Wilcoxon test, FDR ≤ 0.1). **(G-I)** UMAPs of all **(G)**, control **(H**), and disease **(I)** alpha motor neurons with cluster and subtype annotations in (G). **(J)** Point plot showing the proportion of fast-firing alpha motor neurons across control and disease stages (group means ± standard error). Proportions were calculated as the number of fast-firing cells divided by the total number of fast-firing, intermediate, and slow-firing cells. The conditions include n = 3 samples for Control and Mid/End, and n = 2 for Early. An asterisk denotes the p-value (one-sided Welch’s t-test): ** p ≤ 0.01.

To investigate the chromatin accessibility landscapes of motor neurons, we first subclustered and annotated the cholinergic neuron nuclei (Figure S4C-S4D, Figure 3C).^28,32^ We then downsampled the data to obtain equal cell numbers from distinct groups to avoid biasing subsequent analyses towards more abundant groups. At baseline in controls, alpha motor neurons showed the highest number of ATAC-seq fragments and the greatest number of peaks among cholinergic neuron types (Figure 3D–3E), while the fragment size distributions were similar (Figure S4E). In contrast, the transcription start site (TSS) enrichment profiles and scores and the fraction of fragments in peaks were lowest in alpha motor neurons (Figure S4F–S4H). These observations are consistent with a long-observed phenomenon that alpha motor neurons are more euchromatic and transcriptionally active than other cell types.^27,47,48^ We next identified chromatin accessibility changes with disease in vulnerable and resistant motor neurons using the control and mid/end-stage disease data (Figure 3F). The alpha motor neuron epigenome was substantially altered with disease progression in terms of differentially accessible peaks, while resistant pan-gamma (gamma and gamma*) and visceral motor neurons exhibited minimal changes (Figure 3F).

### Chromatin accessibility landscapes of healthy and DAMN alpha motor neurons

To define the chromatin accessibility landscapes of alpha motor neuron subtypes, we subclustered alpha motor neuron nuclei into five groups using the ATAC-seq data (Figure 3G). As with the RNA data, we found disease-dependent, DAMN clusters that were essentially absent in the control samples but present in SOD1-G93A samples (Figure 3H-3I). One DAMN cluster had less extensive gene expression changes (early DAMN), while the other had more extensive changes (late DAMN) and contained alpha motor neurons that were further along in the degeneration/cell death process (Figure 3G, Figure S4I). The remaining three subclusters were well-represented in both control and disease conditions (Figure 3H-3I). Using expression of known marker genes^28,45,46^, we found that two of these clusters corresponded to fast-firing (FF) and slow-firing (SF) alpha motor neurons, respectively (Figure 3G, Figure S4I). The third cluster showed intermediate expression levels of fast- and slow-firing marker genes (Figure 3G, Figure S4I) and may have intermediate electrophysiological properties. These results provide chromatin accessibility landscapes for alpha motor neurons according to DAMN status (non-DAMN, early DAMN, late DAMN) and functional subtype (FF, intermediate, SF).

Consistent with the snRNA-seq data, we saw an increasing representation of DAMNs through disease progression (Figure S4J). Because FF alpha motor neurons are more susceptible to degeneration compared to SF alpha motor neurons in ALS^15–18^, we assessed whether the proportion of FF alpha motor neurons decreased with disease in our mouse data. Indeed, we found a significant decrease in the proportion of FF alpha neurons at mid/end-stage compared to control (Figure 3J), suggesting that FF alpha motor neurons are more likely to enter the DAMN state in disease. We also transferred these subtype labels to the RNA-only alpha motor neuron data and observed concordance between the DAMN clusters identified by the RNA and ATAC modalities (Figure S4K).

### Transcription factors associated with the DAMN state transition

To identify transcription factors (TFs) that may drive the transition from healthy alpha motor neurons to DAMNs, we analyzed the multiome and snRNA-seq data using ArchR^49^ (Figure 4A, Table S1N) and CellOracle^50^ (Figure 4B, Table S1O). ArchR uses chromVAR^51^ to compute TF motif activities (deviation z-scores) for each cell, reflecting the relative accessibility of TF binding sites across cells. Candidate regulators of the DAMN transition are TFs whose motif activities correlate with their own gene expression across alpha motor neurons (Figure 4A, x-axis) and differ across non-DAMN, early DAMN, and late DAMN states (Figure 4A, y-axis, max TF motif delta). CellOracle constructs gene regulatory networks from snATAC-seq and models TF effects using snRNA-seq. It simulates TF knockouts to predict changes in cell state trajectories, quantified for each cell by the perturbation score (ps) and aggregated across cells as the ps sum. Strongly negative ps sum values suggest that the TF may promote DAMN formation (Figure 4B). TFs identified by both methods include C2H2 zinc-finger factors from the Sp/KLF (*Klf6*, *Klf7*) and GLI-Krüppel (*E4f1*) groups as well as bZIP factors from the AP-1 (*Fosl1*, *Jun*, *Jund*), ATF/CREB (*Atf3*, *Atf5*), and PAR-bZIP (*Nfil3*) groups (Figure 4A-4B, Table S1N-S1O). Many of these TFs, along with others, showed significant disease-associated changes in gene expression and corresponding alterations in motif accessibility (Figure 4C-4D). To support these findings, we validated the expression changes of several candidate TFs using MERFISH (Figure 4E, Table S1M).

**Figure 4:**
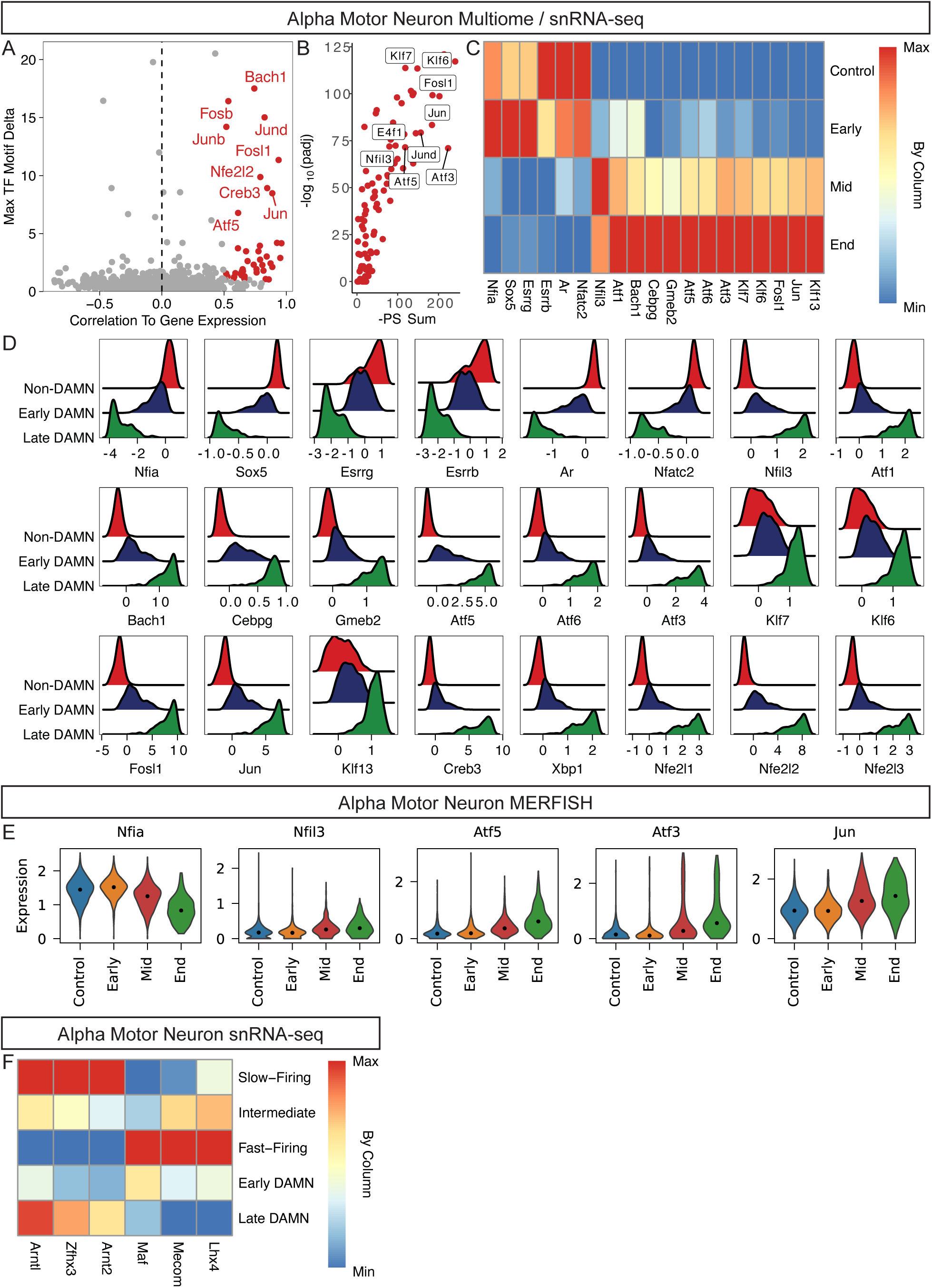
Transcription factor (TF) regulators of the disease-associated motor neuron state. **(A)** Dot plot showing the identification of positive TF regulators of DAMN status (non-, early, late DAMN) with ArchR. TFs with maximum differences in chromVAR deviation z-score in the top quartile of all TFs and a correlation of greater than 0.5 are plotted in red and are candidate positive TF regulators. **(B)** CellOracle *in silico* knockout results of 106 TFs in the non-DAMN to DAMN transition summarized as a volcano plot with the sum of negative perturbation scores on the x-axis and −log_10_(padj) on the y-axis. TFs that were also identified in the ArchR analysis from (A) are labeled. **(C)** Heatmap showing average expression levels of selected differentially expressed TFs throughout disease progression in alpha motor neurons by snRNA-seq. Data are min/max-normalized by column. **(D)** Distributions of chromVAR deviation scores of selected TFs from (A) for non-, early, and late DAMN alpha motor neurons. **(E)** Violin plots showing the log-normalized expression of select differentially expressed genes in alpha motor neurons throughout disease progression by MERFISH (two-sided Mann-Whitney U test between control and end-stage with Benjamini-Hochberg correction; FDR < 0.01). Dots mark the median values. **(F)** Heatmap showing average expression levels of selected differentially expressed TFs in alpha motor neuron subtypes by snRNA-seq. Data are min/max-normalized by column.

Our multiomic profiling enabled high-confidence prediction of TFs that drive DAMN epigenetic remodeling. However, these data cannot on their own determine whether the transcriptional regulators active in DAMNs contribute to pathology, confer protection, or exert both effects simultaneously. To address this, we asked whether any of the TFs nominated by our unbiased analyses have been implicated in either promoting or mitigating ALS pathogenesis. Two of the top DAMN regulators, *Atf3* and *Nfil3* (Figure 4A-4E), demonstrate robust neuroprotection and delayed disease onset when overexpressed in the SOD1-G93A mouse.^52,53^ Additionally, a recent study showed that *CREB3*, another positive regulator of the DAMN state, carries a gain-of-function variant that boosts CREB3 activity and protects against human ALS (Figure 4A, Figure 4D).^54^ Furthermore, several of the other TFs—*Xbp1*, *Nfe2l1*, *Nfe2l2*, and *Nfe2l3* (Figure 4D)— have known roles in promoting protein folding, protein degradation, and recovery from oxidative stress, suggesting they may function as protective regulators. Collectively, our results show that components of the DAMN transcriptional program represent an active attempt to counteract disease pathogenesis and suggest that provoking aspects of the DAMN signature via specific TFs is protective in the context of SOD1 ALS.

### Transcription factors associated with vulnerable and resistant skeletal motor neurons

Beyond the DAMN transcription factors (TFs), we hypothesize that TFs may also be responsible for gene regulatory networks that determine a motor neuron’s intrinsic differential vulnerability to disease (e.g., alpha vs. pan-gamma skeletal motor neurons or FF vs. SF alpha motor neurons). Thus, we sought to identify TFs that define selectively vulnerable and resistant motor neuron populations at baseline in the adult spinal cord, building on prior work characterizing TFs in motor neuron development.^55–59^ To do so, we analyzed the control multiome and snRNA-seq data using ArchR^49^ and CellOracle^50^, as described above, focusing on skeletal motor neuron subtypes (alpha, gamma*, gamma) and alpha motor neuron subtypes (FF, intermediate, SF). On top of known motor neuron subtype-specific TFs (e.g. *Esrrg*, *Esrrb*)^28,57,60^, we identified additional TFs that are linked to specific motor neuron subtypes (Figure S5A-S5G, Table S1P-S1T). Interestingly, we found five TFs (*Esrrg*, *Esrrb*, *Mitf*, *Arntl*, and *Arnt2*) that are important for gene regulatory networks of pan-gamma compared to alpha motor neurons as well as SF compared to FF alpha motor neurons (Figure S5A-S5F, Table S1P-S1T), providing a set of TFs that are shared among resistant motor neurons. We also identified two shared TFs from the ArchR and CellOracle analyses of alpha motor neuron subtypes, *Maf* and *Arnt2*, which are associated with FF and SF alpha motor neurons, respectively (Figure S5D-S5G, Table S1R-S1T).

To further nominate alpha motor neuron subtype-specific TFs that may be relevant to disease, we looked for those that also had altered expression between early DAMN and late DAMN cells. We found that *Arnt2*, *Zfhx3*, and *Arntl* were more active in SF compared to FF alpha motor neurons and had increased expression with DAMN progression (Figure 4F, Table S1Q-S1U), suggesting a potential protective role of these TFs in disease (because SF motor neurons are relatively resistant to degeneration and TF networks upregulated in DAMNs are likely a protective response). In contrast, *Maf*, *Mecom*, and *Lhx4* showed the opposite pattern; they were more active in FF compared to SF alpha motor neurons and had decreased expression with DAMN progression (Figure 4F, Table S1Q-S1U), suggesting a potential deleterious role of these TFs in disease. Interestingly, Mecom is necessary for the specification of fast motor neurons in zebrafish^61^, and Maf regulates fast type IIb myofiber determination in mice^62,63^, raising the possibility that shared transcriptional programs may contribute to fast identity across the motor unit. Overall, we identified TFs associated with vulnerable and resistant motor neurons, which may be responsible for specifying motor neuron subtypes in health and influencing disease state (DAMN vs. non-DAMN) during neurodegeneration.

### Glia become reactive near motor neurons early in disease and then become widespread

In addition to cell-autonomous mechanisms of motor neuron degeneration, non-cell autonomous mechanisms contribute to ALS.^64–68^ Toxic properties of glial cells (astrocytes, oligodendrocytes, and microglia) collaborate to drive disease progression.^66^ However, it remains unclear if these glial-specific changes occur independently of motor neurons or if they are triggered as a response to signals from motor neurons. To investigate the relationship between glia and motor neurons in disease, we used spatial transcriptomics to identify changes in cell types that are nearest neighbors to skeletal motor neurons. We found a striking increase in microglia/macrophages near DAMN alpha motor neurons compared to non-DAMNs (Figure 5A, Table S1V) and a similar increase in microglia/macrophages near all skeletal motor neuron subtypes in disease conditions compared to the control condition (Figure S6A, Table S1W). In contrast, we did not observe such increases for other cell classes near DAMNs (Figure 5A, Table S1V). These findings prompted us to assess changes in cell type abundance with disease relative to the number of interneurons. We found a large, ∼10-fold increase in the relative abundance of microglia/macrophages with disease, echoing previous observations,^69^ along with more modest increases in astrocyte and oligodendrocyte abundance (Figure 5B, S6B). These changes coincide with a significant decrease in the relative abundance of alpha motor neurons but not gamma or gamma* motor neurons with disease (Figure S6B). We confirmed that the large increase in microglia/macrophages was not caused by preferential removal of control cells during data pre-processing (Figure S5C), supporting a genuine rise with disease.

**Figure 5:**
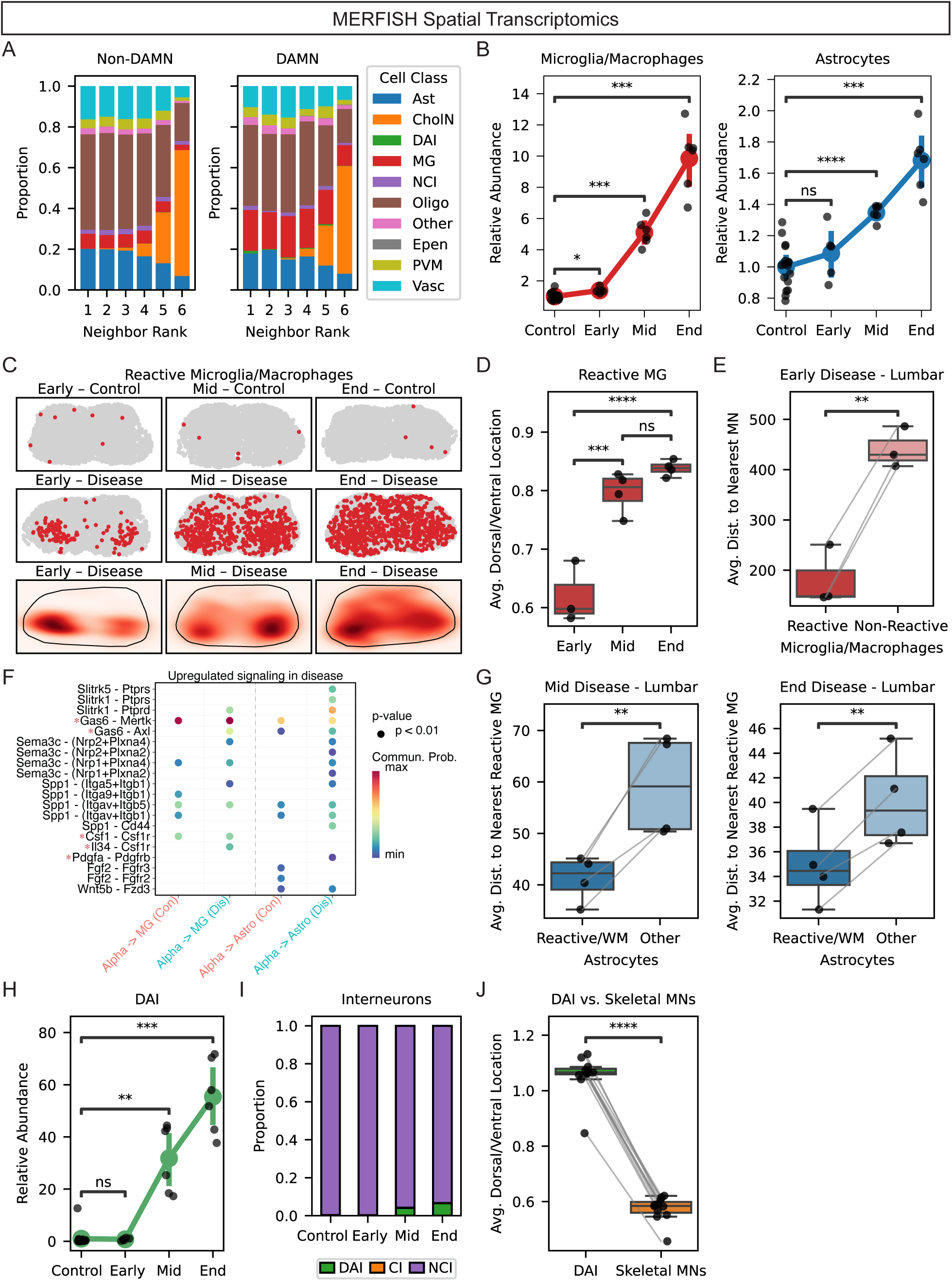
Spatial and compositional dynamics of spinal cord cell types with disease in the SOD1-G93A mouse. **(A)** Stacked bar plots showing the cell class composition of the six nearest neighbors to Non-DAMN and DAMN alpha motor neurons (Ast: Astrocytes, CholN: Cholinergic Neurons, DAI: Disease-Associated Interneurons, MG: Microglia/Macrophages, NCI: Non-Cholinergic Interneurons, Oligo: Oligodendrocytes, Epen: Putative Ependymal Cells, PVM: Putative Perivascular/Meningeal Cells, Vasc: Putative Vascular Cells). Microglia/macrophages were significantly enriched at all neighbor positions for DAMNs compared to Non-DAMNs (Fisher’s Exact Test, Bonferroni-adjusted padj ≤ 0.01). **(B)** Strip and point plots showing the abundance of microglia/macrophages (left) and astrocytes (right) relative to interneurons across control and disease conditions. Each point represents the average value from tissue sections of a MERFISH experiment (n ≥ 5 experiments per condition). Values are scaled to the mean relative abundance in control samples. Colored points and lines indicate cell type means ± 95% confidence intervals. Asterisks indicate significance relative to control (one-sided Welch’s t-test, Bonferroni-adjusted): n.s. = not significant; * padj ≤ 0.05; ** padj ≤ 0.01; *** padj ≤ 0.001; **** padj ≤ 0.0001. **(C)** Spatial plots showing the distribution of reactive microglia/macrophages across control and disease conditions. Top two rows: dots represent individual cells from MERFISH sections (reactive microglia/macrophages: red, other cells: gray). Bottom row: kernel density plots with tissue boundary outlines. **(D)** Box and strip plots showing the average dorsal-ventral position of reactive microglia/macrophages in the lumbar spinal cord across disease stages. Values < 1 indicate positions more ventral, and values > 1 indicate positions more dorsal, than the median cell location within each section. Each point represents the average value from tissue sections of a MERFISH experiment (n ≥ 3 experiments per stage). Asterisks denote adjusted p-values (one-way ANOVA followed by Tukey’s HSD test): n.s. = not significant; *** p ≤ 0.001; **** p ≤ 0.0001. **(E)** Box and strip plots comparing the average distance in μm from reactive and non-reactive microglia/macrophages to the nearest skeletal motor neuron in the lumbar spinal cord during early-stage disease. Each point represents the average value from tissue sections of a MERFISH experiment (n = 3 experiments). Lines connect paired measurements from the same experiment. Asterisks denote p-values (one-sided paired t-test): n.s. = not significant; ** p ≤ 0.01. **(F)** Bubble plot of ligand-receptor interactions upregulated in disease with alpha motor neurons as the sender. Bubble size indicates statistical significance, and color reflects communication probability. Red stars indicate ligand-receptor interactions where the ligand is significantly upregulated in alpha motor neurons at end-stage by snRNA-seq. **(G)** Box and strip plots comparing the average distance in μm from reactive/white matter and other astrocytes to the nearest reactive microglia/macrophage in the lumbar spinal cord during mid-stage (left) or end-stage (right) disease. Each point represents the average value from tissue sections of a MERFISH experiment (n = 4 experiments). Lines connect paired measurements from the same experiment. Asterisks denote p-values (one-sided paired t-test): ** p ≤ 0.01. **(H)** Strip and point plots showing the abundance of disease-associated interneurons (DAI) relative to other interneurons across control and disease conditions. Each point represents the average value from tissue sections of a MERFISH experiment (n ≥ 5 experiments per condition). Values are scaled to the mean relative abundance in control samples. Colored points and lines indicate cell type means ± 95% confidence intervals. Asterisks indicate significance relative to control (one-sided Welch’s t-test, Bonferroni-adjusted): n.s. = not significant; ** padj ≤ 0.01; *** padj ≤ 0.001. **(I)** Stacked bar plots showing the proportions of interneuron subtypes across control and disease conditions (DAI: Disease-Associated Interneurons, CI: Cholinergic Interneurons, NCI: Non-Cholinergic Interneurons). **(J)** Box and strip plots showing the average dorsal-ventral position of disease-associated interneurons (DAI) and skeletal motor neurons in the lumbar spinal cord during disease (early, mid, and end-stage). Values < 1 indicate positions more ventral, and values > 1 indicate positions more dorsal than the median cell location within each tissue section. Each point represents the average value from tissue sections of a MERFISH experiment (n = 11 experiments). Lines connect paired measurements from the same experiment. Asterisks denote p-values (one-sided paired t-test): n.s. = not significant; **** p ≤ 0.0001.

To further assess changes in glia with disease, we identified differentially expressed genes with the snRNA-seq data and found upregulation of reactive astrocyte (Table S1X) and disease-associated microglia genes (Table S1Y) as well as downregulation of homeostatic microglia genes (Table S1T), among other changes.^70–72^ In the MERFISH data, we used *Apoe* and *Gfap* expression to classify microglia and astrocytes as reactive/white matter or non-reactive. Consistent with prior work, we found that the proportion of glia in a reactive state increases with disease progression^73–75^ along with an increase in the proportion of DAMNs (Figure S6D-S5F). Strikingly, we found that reactive microglia/macrophages were spatially located near motor neurons early in disease and became more widespread throughout the spinal cord as disease progressed (Figure 5C-5E).^74,76^ We also found that reactive/white matter astrocytes were significantly closer to reactive microglia/macrophages than other astrocytes were at mid and end-stage disease (Figure 5F-5G), consistent with the known role of reactive microglia in activating astrocytes.^68,71^

The emergence of reactive and proliferating microglia near motor neurons in the ventral horn preceded widespread alpha motor neuron death (Figure 5B-5E, Figure S6B). Thus, we posited that there may be a signal emanating from DAMNs that activates local microglia. To prioritize well-established monocytic signaling pathways, we queried the ChatGPT o3 model to provide candidate signaling ligands that are upregulated in DAMNs and could promote microglial reactivity/proliferation. The analysis highlighted *Csf1*, *Il34*, and *Lgals3* as candidates (Table S1E). We also used CellChat^77^ with the snRNA-seq data to identify disease-associated ligand-receptor pairs among alpha motor neurons, microglia/macrophages, and astrocytes based on differential expression analysis. CellChat predicted baseline CSF1 – CSF1R signaling from alpha motor neurons to microglia/macrophages and increased CSF1 – CSF1R signaling from all three cell types to microglia/macrophages in disease (Figure 5F, Figure S6G-S6H). It also predicted IL-34 – CSF1R signaling specifically from alpha motor neurons to microglia/macrophages in disease, along with additional disease-associated ligand-receptor signaling events (Figure 5F, Figure S6G-S6H). Prior studies show that CSF1R activation causes a proliferative phenotype^78–80^ and that damaged peripheral sensory^81,82^ and motor neurons^82,83^ upregulate CSF1, which directly induces microglial activation. Together, our results show that *Il34* and *Csf1* are both induced as a part of the DAMN signature, coincident with increased microglial proliferation and reactivity (Figure 5B-5C, Table S1E). Notably, the literature suggests that CSF1 and IL-34 can have opposing effects on microglia, supporting the idea that the DAMN signature may have both adaptive and maladaptive consequences.

The changes in reactive glia coincided with an increased abundance of disease-associated interneurons (DAI) relative to other interneurons at mid and end stages (Figure 5H). DAI were also observed in our snRNA-seq and multiome sequencing data (Figure 1B, Figure 3B), consistent with reported dysfunction of interneurons in this model.^84–86^ Among all interneurons, DAI increased in abundance with disease progression up to ∼7% at end-stage (Figure 5I), while ∼56% of alpha motor neurons were in the DAMN state at end-stage (Figure S6G). DAI were, on average, located more dorsally in the spinal cord compared to motor neurons (Figure 5J) and may contribute to or result from the increase in reactive glia in more dorsal locations with disease progression. Future work is required to elucidate the interactions among reactive glia, DAMNs, and DAI, but these findings raise the possibility that early glial reactivity may be influenced by signals from nearby motor neurons, despite broad mutant SOD1 expression.

### Molecular changes in motor neurons connect the SOD1-G93A mouse to human ALS

In human ALS, only about 2% of cases are caused by *SOD1* mutations and associated SOD1 pathology.^3,4^ Nearly all other cases are instead accompanied by nuclear depletion and cytoplasmic aggregation of the DNA/RNA-binding protein TDP-43.^87^ Thus, we wanted to investigate whether the pathways involved in the DAMN state are specific to *SOD1* mutations or are connected to human ALS genetics more broadly. We took advantage of genome-wide association studies (GWAS)—a population genetics technique that identifies common genetic variants associated with human traits of interest, such as ALS—to determine if the regions that are differentially accessible with disease in alpha motor neurons (Figure 6A) are enriched for disease-associated single nucleotide polymorphisms (SNPs). We converted ∼40% of these disease peaks from mouse to the human genome with high confidence, and we used these human genomic regions along with GWAS summary statistics to perform partitioned linkage disequilibrium (LD) score regression analyses (Figure 6B).^88,89^ This allowed us to test whether genomic regions with differential accessibility in disease disproportionately contribute to the heritability of ALS (Figure 6B).

**Figure 6:**
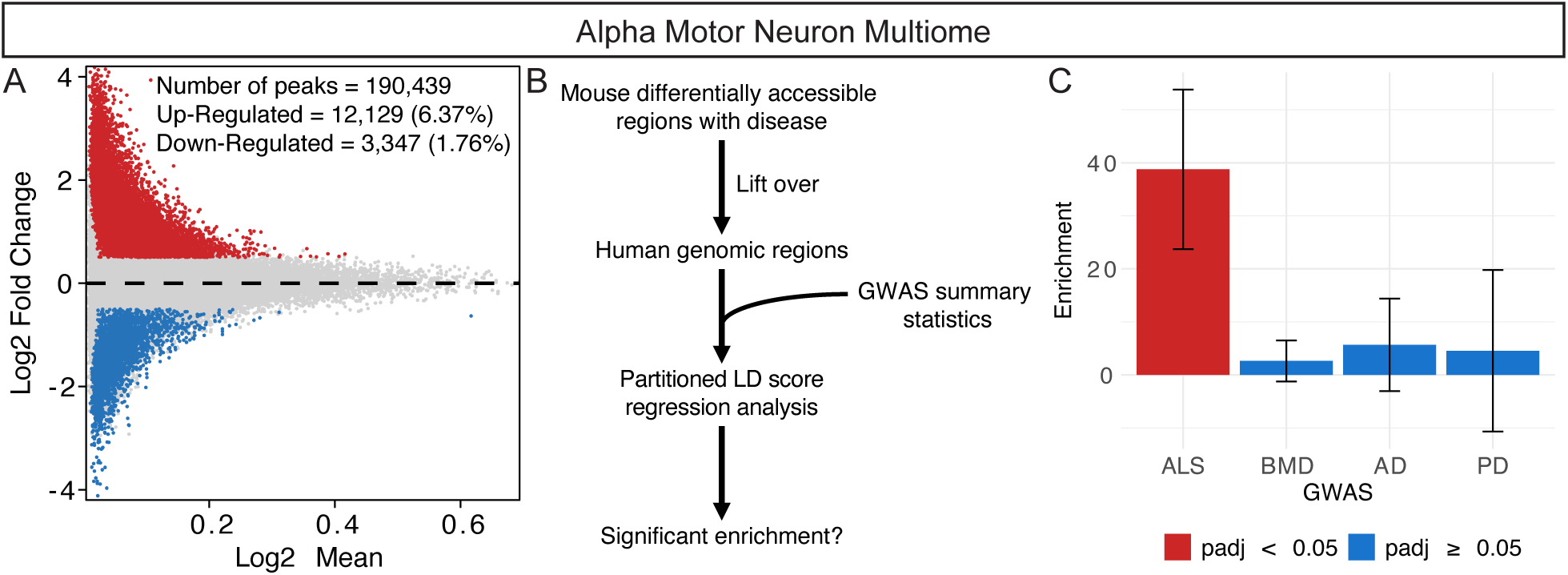
Linking chromatin accessibility changes in SOD1-G93A motor neurons to human ALS. **(A)** MA plot showing differentially accessible peaks between mid/end-stage and control alpha motor neurons. Each point represents a peak, with the x-axis indicating the log₂ mean accessibility across all control and mid/end-stage alpha motor neurons and the y-axis showing the log₂ fold change. Significant peaks (Wilcoxon test; FDR ≤ 0.05, |log₂FC| ≥ 0.5) are highlighted in red (upregulated in disease) and blue (downregulated in disease). **(B)** Schematic of the analysis pipeline used to assess the relevance of chromatin accessibility changes in mouse alpha motor neurons to human disease. Differentially accessible regions with disease from mouse alpha motor neurons were lifted over to the human genome. Partitioned LD score regression analysis was then used to test whether these regions were enriched for SNP heritability using GWAS summary statistics. **(C)** Bar plot of heritability enrichment scores from partitioned LD score regression across four traits using differentially accessible peaks identified in (A) (FDR < 0.05). Traits include ALS (Amyotrophic Lateral Sclerosis), BMD (Bone Mineral Density), AD (Alzheimer’s Disease), and PD (Parkinson’s Disease). Bars show enrichment estimates with error bars (± standard error). Bars are colored by statistical significance (red: padj < 0.05; blue: padj ≥ 0.05).

Our results demonstrate significant heritability enrichment when using GWAS summary statistics from human ALS^90^ (Figure 6C), but no such enrichments when using those from a non-brain-related trait (bone mineral density abnormality: BMD) or from other neurodegenerative disorders (Alzheimer’s disease: AD and Parkinson’s disease: PD) (Figure 6C).^91–93^ The ALS GWAS has fewer samples than the other studies do (152,268 for ALS, 426,824 for BMD, 788,989 for AD, and 2,525,897 for PD)^90–93^, suggesting that this result is not dependent on the sample size of the GWAS. To identify genes mediating the ALS heritability enrichment, we performed Hi-C coupled Multi-marker Analysis of GenoMic Annotation (H-MAGMA) with the most recent ALS GWAS^90^ and intersected the output with genes located in the differentially accessible regions (Figure S7A, Table S1Z).^94,95^ A handful of the prioritized genes are significantly differentially expressed with disease in SOD1-G93A mouse alpha motor neurons and may contribute to disease pathogenesis (Figure S7B). Collectively, our results suggest that despite the upstream triggers of degeneration being different (mutant SOD1 transgene in mouse vs. various genetic and pathological triggers in human), aspects of the genetic basis of motor neuron degeneration are shared between the SOD1-G93A mouse model and human ALS.

## Discussion

By employing longitudinal single-nucleus RNA/ATAC-seq and spatial transcriptomics throughout disease progression in the SOD1-G93A mouse model of ALS, we uncovered extensive molecular changes in a subset of vulnerable alpha motor neurons. Alpha motor neurons with these gene expression and chromatin accessibility changes have transitioned from healthy cells into a new cell state, which we have named ‘disease-associated motor neurons’ (DAMNs). Motor neurons that are more resistant to degeneration (e.g., pan-gamma and visceral motor neurons) exhibited a relatively minor response to disease, but vulnerable alpha motor neurons (especially FFs) readily became DAMNs, highlighting the importance of studying motor neuron subtypes individually rather than as a single population.

Just as SOD1-G93A mice develop hindlimb motor impairment before the forelimbs are affected^7^, individual alpha motor neurons do not transition to the DAMN state all at once—we observed a small number of DAMNs before overt behavioral phenotypes were apparent at the organismal level as well as many non-DAMN alpha motor neurons that persisted even at later stages of disease. These findings argue that perhaps we should frame disease onset from a cellular perspective, based on gene expression and functional properties, as well as from an organismal level, based on behavior and survival. It also suggests that even though it may be too late to protect DAMNs, the remaining non-DAMN alpha motor neurons could still be viable targets for therapeutic intervention. We also uncovered previously unrecognized temporal dynamics of transcriptional changes in alpha motor neurons. Among these, early alterations may provide insight into the initial steps of disease pathogenesis, and modulating such changes could help slow motor neuron degeneration.

The transcriptional changes distinguishing DAMNs from healthy alpha motor neurons arise, at least in part, from specific transcription factors that we have nominated based on our ATAC and RNA sequencing data. We cannot rule out the possibility that some transcription factor changes associated with the DAMN state may promote disease pathogenesis, but we hypothesize that many of these changes represent protective responses that occur too late to prevent motor neuron death. This hypothesis is supported by previous findings showing that increased activity of transcription factors associated with the DAMN state are protective in the context of disease.^52–54^ We additionally identified transcription factors linked to vulnerable and resistant motor neuron subtypes, a subset of which are also perturbed in disease. Future studies aimed at further understanding the regulatory networks governing motor neuron subtype-selective vulnerability and the DAMN state transition will reveal novel therapeutic targets for ALS.

It will be important to comprehensively determine the extent to which the DAMN state overlaps between mouse models and human ALS in future work. Nevertheless, prior studies examining individual components of the DAMN signature in postmortem samples suggest that it is at least partially conserved. For example, human orthologs of DAMN-upregulated *Gap43*, *Ngf,* and *Jun*^96–98^ all show increases in RNA levels or encoded protein products in postmortem sporadic ALS spinal cords, while human orthologs of DAMN-downregulated *Gria2* and *Gria4* show dramatic downregulation in patient motor neurons from sporadic and *SOD1* ALS cases^99^.

In addition to cell-intrinsic programs, we observed a striking increase in reactive microglia/macrophages near motor neurons early in disease, followed by widespread glial reactivity at later stages. This early proximity was unexpected, given that the mutant SOD1 transgene is expressed in all cells and could have driven glial activation through purely cell-autonomous mechanisms. Instead, the pattern suggests that local signals from motor neurons may contribute to early glial activation. We identified potential candidate ligands that are upregulated in DAMNs (e.g., *Csf1* and *Il34*), which could be conserved in human patients. CSF1 levels increase in sporadic and SOD1 patient spinal cords and peripheral nerves^100,101^ and in peripheral nerves in the SOD1 rat model.^100^ IL-34, which is required for the differentiation of microglia from monocytes and is required for maintaining expanding homeostatic (non-reactive) microglia, has not specifically been studied in the context of motor neurons. Because both are direct ligands of CSF1R, this finding argues against broad inhibition of CSF1R in favor of a more targeted approach to selectively inhibit CSF1 expression in motor neurons, perhaps via genetic knockout or antisense oligonucleotide-based therapies. While it merits further interrogation, we propose a preliminary model in which the cascade of glial activation begins with CSF1 and IL-34 stimulating microglial proliferation but eventually CSF1 tips microglia into reactivity, causing microglia to (a) increase autocrine CSF1 signaling and (b) secrete IL-1α, TNFα, and C1q that activate astrocytes^68,71^, culminating in increased astrocyte-secreted toxic factors that kill neurons^102^.

The underlying pathology of ALS patients with *SOD1* mutations is distinct from that of patients with other mutations or sporadic disease, leading many to speculate that SOD1 ALS may be a different subtype of the disease.^1^ However, SOD1 ALS patients are clinically indistinguishable from sporadic ALS patients, suggesting that there may be convergent downstream molecular pathways that drive motor neuron degeneration. We found that the human counterparts of regions with differential accessibility in SOD1-G93A alpha motor neurons are enriched for ALS-associated SNPs. This finding shows that genetic variations that combine to confer risk for ALS in humans impact the same pathways that contribute to degeneration in the SOD1-G93A mouse. Thus, the SOD1 mouse can be a powerful model to study fundamental aspects of motor neuron degeneration and to develop and test therapies aimed at increasing motor neuron resilience and preventing the pathological aspects of the DAMN state.

By defining the transcriptional and epigenomic landscapes of vulnerable and resilient motor neuron populations throughout neurodegeneration, we lay the groundwork for future investigations aimed at rescuing disease phenotypes and developing therapeutic avenues. Continued exploration of the interplay between genetic factors, transcriptional regulation, and motor neuron subtype-specific responses will be essential for advancing our understanding of this devastating disease.

## Acknowledgments

This work was supported by NIH grants to A.D.G. (R35NS137159, U54NS123743, R01AG064690), W.J.G. (R01NS128028), J.E.R. (R01AG075802, RF1NS128800), and N.S. (RM1HG010461); the Robert Packard Center for ALS Research at Johns Hopkins (A.D.G.); Target ALS (A.D.G and M.Y.); the Phil and Penny Knight Initiative for Brain Resilience (A.D.G. and Y.Z.); the National Science Foundation Graduate Research Fellowship (DGE-1656518) (O.G.); a Stanford Graduate Fellowship (O.G.); the Blavatnik Family Foundation (J.A.B.); the American Neuromuscular Foundation (M.Y.); and the Larry L. Hillblom Foundation (Y.Z.). A.D.G. and W.J.G. are Chan Zuckerberg Biohub Investigators. Sorting was performed on an instrument in the Shared FACS Facility obtained using NIH S10 Shared Instrument Grant S10RR025518.

## Competing Interests

A.D.G. is a scientific founder of Maze Therapeutics, Trace Neuroscience, and Lyterian Therapeutics. W.J.G. has affiliations with 10x Genomics (consultant), Guardant Health (consultant) and Protillion Biosciences (co-founder and consultant).

### Supplemental Information

- **Table S1A:** Differential expression analysis - Alpha motor neurons (end-stage vs. control)
- **Table S1B:** Differential expression analysis - Gamma* motor neurons (end-stage vs. control)
- **Table S1C:** Differential expression analysis - Gamma motor neurons (end-stage vs. control)
- **Table S1D:** Shared upregulated and downregulated genes in motor neurons with disease and sciatic nerve crush
- **Table S1E:** Differential expression analysis - DAMNs vs. non-DAMNs
- **Table S1F:** Differential expression analysis - pre-DAMN fast-firing alpha motor neurons (mid/end-stage vs. control)
- **Table S1G:** Differential expression analysis - pre-DAMN slow-firing alpha motor neurons (mid/end-stage vs. control)
- **Table S1H:** Upregulated GO biological processes in pre-DAMN fast-firing alpha motor neurons
- **Table S1I:** Downregulated GO biological processes in pre-DAMN fast-firing alpha motor neurons
- **Table S1J:** Upregulated GO biological processes in DAMNs
- **Table S1K:** Downregulated GO biological processes in DAMNs
- **Table S1L:** List of genes in the MERFISH panel
- **Table S1M:** Differential expression analysis - MERFISH Alpha motor neurons (mid-stage and end-stage vs. control)
- **Table S1N:** Positive transcription factor regulators of DAMNs
- **Table S1O:** CellOracle *in silico* transcription factor knockout results for DAMNs
- **Table S1P:** Positive transcription factor regulators of skeletal motor neuron subtypes (Alpha, Gamma*, Gamma)
- **Table S1Q:** Differential expression analysis - slow-firing vs. fast-firing alpha motor neurons
- **Table S1R:** Positive transcription factor regulators of alpha motor neuron subtypes (fast-firing, intermediate, slow-firing)
- **Table S1S:** CellOracle *in silico* transcription factor knockout results for fast-firing alpha motor neurons
- **Table S1T:** CellOracle *in silico* transcription factor knockout results for slow-firing alpha motor neurons
- **Table S1U:** Differential expression analysis - early DAMNs vs. late DAMNs
- **Table S1V:** DAMN nearest-neighbor analysis
- **Table S1W:** Skeletal motor neuron nearest-neighbor analysis
- **Table S1X:** Differential expression analysis - Astrocytes (end-stage vs. control)
- **Table S1Y:** Differential expression analysis - Microglia/Macrophages (end-stage vs. control)
- **Table S1Z:** H-MAGMA results

## Methods

### Animals

All procedures involving mice were performed in accordance with a protocol approved by the Administrative Panel of Laboratory Animal Care of Stanford University (protocol no. 30643). For snRNA-seq and multiome sequencing experiments, ChAT-IRES-Cre mice (B6;129S6-*Chattm2(cre)Lowl*/J; Jax Strain #: 006410) were crossed with ROSAnT-nG mice (B6;129S6-*Gt(ROSA)26Sortm1(CAG-tdTomato*,-EGFP*)Ees*/J; Jax Strain #: 023035) to generate mice homozygous for both alleles. Resulting female mice were crossed with male hemizygous SOD1-G93A mice (B6SJL-Tg(SOD1*G93A)1Gur/J; Jax Strain #: 002726). Male and female progeny that carried the SOD1-G93A transgene were used for the early-, mid-, and end-stage SOD1 experiments, and progeny that lacked the SOD1-G93A transgene were used for age-matched control experiments. For MERFISH experiments, male SOD1-G93A mice (B6SJL-Tg(SOD1*G93A)1Gur/J; Jax Strain #: 002726) and age-matched wild-type male mice (B6SJLF1/J; Jax Strain #: 100012) were used. All mice were housed with food and water available *ad libitum* in a 12-h light/dark environment at ambient temperature and humidity. Symptomatic mice carrying the SOD1-G93A transgene were given HydroGel (ClearH2O) and food on the cage floor. All animals were euthanized at or before the humane euthanasia point, which was defined as the inability of a mouse to right itself within 20 s after being placed on its back or side.

### Nuclei Isolation and Fluorescence-Activated Nuclei Sorting (FANS)

Nuclei isolation and sorting was adapted from Blum et al. (2021), and buffer recipes are found in Blum et al. (2021). Twenty-four snRNA-seq experiments and five multiome (paired snATAC/snRNA-seq) experiments were performed. For each experiment, several mice (∼3-12) were euthanized with CO2. The spinal cords were rapidly hydraulically extruded using a PBS-filled syringe with a blunt 21 G needle. Two spinal cords were homogenized at a time in a 2 mL Dounce homogenizer (Sigma-Aldrich, D8938-1SET) containing 2 mL of nuclei extraction buffer. The spinal cords were homogenized with ten strokes of pestle A followed by five strokes of pestle B. The homogenate from all cords was transferred to a 50 mL round-bottom centrifugation tube (Thermo Scientific, 3118-0050PK), and 8 mL of nuclei spin buffer 1 was added. Then, 5 mL of nuclei spin buffer 2 was layered gently underneath the homogenate solution. The nuclei were pelleted through the gradient in a swinging-bucket centrifuge (3200xg, 4°C, 15 minutes). The supernatant was rapidly discarded, and the nuclei were resuspended in 5 mL of nuclei spin buffer 1. Next, 5 mL of nuclei buffer 3 was gently layered underneath the nuclei solution. The nuclei were pelleted through the gradient in a swinging-bucket centrifuge (3200xg, 4°C, 15 minutes). The supernatant was rapidly discarded, and the pellet was resuspended in loading buffer containing DAPI.

FANS was performed on a BD Biosciences FACSAria II flow cytometer using the 70 μm nozzle. Single nuclei were gated using DAPI and side scatter measurements to exclude debris and multiplets. Among the single nuclei, EGFP+/tdTomato- and tdTomato+/EGFP-nuclei were identified using a two-dimensional scatterplot. Nuclei were sorted such that ∼40% were EGFP+/tdTomato- and ∼60% tdTomato+/EGFP-for the snRNA-seq experiments, and ∼55% were EGFP+/tdTomato- and ∼45% tdTomato+/EGFP-for the multiome sequencing experiments.

### 10x Genomics snRNA-seq and Multiome Sequencing

Following nuclei sorting, either snRNA-seq (Chromium Single Cell 3′ Reagent Kit v3, 10x Genomics) or multiome ATAC + gene expression profiling (Chromium Single Cell Multiome ATAC + Gene Expression v1, 10x Genomics) was performed according to manufacturer protocols. Libraries were sequenced on an Illumina NextSeq 550 or NovaSeq 6000 using run parameters specified by 10x Genomics.

### snRNA-seq: Data Pre-Processing and Ambient RNA Removal

Sequencing reads were demultiplexed and aligned to a custom mouse pre-mRNA reference transcriptome containing mutant hSOD1 (10x Genomics) using the cellranger mkfastq and cellranger count pipelines (Cell Ranger v3.1.0, 10x Genomics). Subsequent analyses were performed in R (v4.1.1) using the Seurat package (v4.0.4).^103^ A Seurat object was made for each sample using the filtered count matrix which excludes putative empty droplets. Each Seurat object was processed to obtain clusters (NormalizeData, FindVariableFeatures, ScaleData, RunPCA, FindNeighbors with dims = 1:20, FindClusters with resolution = 0.5), and these clusters were used with SoupX (v1.6.2)^104^ to remove ambient RNA contamination (SoupChannel, setClusters, autoEstCont). The SoupX-adjusted counts were rounded to the nearest integer. Mitochondrial reads were then removed and stage information (“ctl”, “sod.early”, “sod.mid”, “sod.end”) was added to the metadata of each object. Some samples were derived from a mixture of both SOD1-G93A mice of a given stage and control mice. For these samples, each RNA profile was assigned to the appropriate condition using the presence or absence of hSOD1 counts.

### snRNA-seq: Integration, Quality Control, and Clustering

The Seurat objects were integrated using Seurat’s reference-based integration workflow. First, the Seurat objects were log-normalized (NormalizeData), and variable features were identified for each object individually (FindVariableFeatures with selection.method = “vst” and features = 2000). Features that were repeatedly variable across datasets were selected for integration (SelectIntegrationFeatures). The sample “210727_nova_CZI/sod1_mn_nuclei_2_9”, which was derived from control and early-stage SOD1-G93A mice, was selected as the reference. The reference was used to find integration anchors (FindIntegrationAnchors), and then the integrated object was created (IntegrateData).

Clustering and visualization were performed using the integrated Seurat object (DefaultAssay set to “integrated”, ScaleData, RunPCA, RunUMAP with reduction = “pca” and dims = 1:20, FindNeighbors with reduction = “pca” and dims = 1:20, FindClusters with resolution = 2). scDblFinder (v1.8.0)^105^ was to identify doublets. Clusters with >40% of the RNA profiles classified as “doublet” by scDblFinder were removed, and any remaining RNA profiles classified as “doublet” were also removed from the Seurat object. Next, clusters with high expression of marker genes that are known to be mutually exclusive were removed to eliminate additional doublet clusters. The remaining RNA profiles were re-clustered (DefaultAssay set to “integrated”, FindNeighbors with reduction = “pca”, and dims = 1:20, FindClusters with resolution = 0.8).

To remove clusters with a low proportion of intron-containing reads, we analyzed a representative end-stage sample (“3nextseqs_11_18/3nextseq_run_11_8_endpoint_female”). Raw count matrices were generated using cellranger count (Cell Ranger v7.1.0, 10x Genomics) with the mm10-2020-A reference transcriptome, with the --include-introns flag set to true and false. The resulting matrices were processed using Read10X and CreateSeuratObject. For each barcode, the proportion of intronic reads (prop_intronic) was calculated as the difference between total (exonic + intronic) and exonic-only UMI counts, divided by the total. Clusters with a median prop_intronic below 0.4 were excluded from further analysis. The final clusters were manually annotated in the cell_class metadata column using the expression of known marker genes.^28,30^

### snRNA-seq: Cholinergic Neuron and Alpha Motor Neuron Subclustering

A Seurat object containing data from cholinergic neurons was created using the nuclei annotated as “Cholinergic Neurons” from the above analysis. Variable features were identified (DefaultAssay set to “RNA”, FindVariableFeatures), and subclustering was performed (DefaultAssay set to “integrated”, ScaleData, RunPCA, RunUMAP with reduction = “pca” and dims = 1:13, FindNeighbors with reduction = “pca” and dims = 1:13, FindClusters with resolution = 0.4). The resulting clusters were manually annotated in the cholingergic_type metadata column using the expression of known marker genes from mouse cholinergic neurons.^28,32^ Another Seurat object containing data from alpha motor neurons was created using the nuclei annotated as “Alpha MNs” from the previous analysis of cholinergic neurons. Variable features were identified (DefaultAssay set to “RNA”, FindVariableFeatures), and subclustering was performed (DefaultAssay set to “integrated”, ScaleData, RunPCA, RunUMAP with reduction = “pca” and dims = 1:12, FindNeighbors with reduction = “pca” and dims = 1:12, FindClusters with resolution = 0.7).

### snRNA-seq: Differential Expression and Gene Ontology (GO) Enrichment Analyses

DESeq2 (v1.32.0)^106^ was used to perform pseudobulk differential expression analyses. For each analysis, SoupX-adjusted counts rounded to the nearest integer were aggregated by gene across all nuclei of a given type for each experiment/biological replicate. All pseudobulk replicates contained data from at least 30 nuclei. The resulting genes-by-replicates count matrix was used to create a data frame of sample information, which included the condition of interest for the differential expression analysis (ex: stage information or cell type, see custom code). The count matrix and sample information were used as input into DESeqDataSetFromMatrix with design set to the condition of interest. The resulting object was used as input into the DESeq function (minReplicatesForReplace = Inf) to perform differential expression analysis using the Wald test to compute statistical significance and the Benjamini-Hochberg method to correct for testing of multiple hypotheses. The differential expression results were extracted, and genes with an adjusted p value of NA were removed. EnrichR (https://maayanlab.cloud/Enrichr/#)^107,108^ was used to perform GO enrichment analysis. Significantly differentially expressed genes (padj < 0.01) that were upregulated or downregulated were used as the input gene set, and the background gene set was all genes from the differential expression analysis that had a numerical adjusted p value. Results from the GO Biological Process 2025 analysis were used. These analyses used the Fisher exact test to compute statistical significance and the Benjamini-Hochberg method to correct for testing of multiple hypotheses. An adjusted p value of < 0.1 was used to determine significantly enriched GO biological processes.

### Multiome Sequencing: Data Pre-Processing

Sequencing reads for five samples were demultiplexed and aligned to a custom mouse mm10-arc reference, which allows mutant hSOD1 gene expression to be detected (10x Genomics), using the cellranger-arc mkfastq and cellranger-arc count pipelines (Cell Ranger ARC v2.0.1, 10x Genomics). Subsequent analyses were performed in R (v4.1.1) using the ArchR package (v1.0.2)^49^ unless otherwise noted. Arrow files were generated from ATAC fragment files (createArrowFiles, default parameters) and used to construct an ArchRProject. To incorporate gene expression data, filtered gene-barcode matrices from each sample were imported (import10xFeatureMatrix) and added to the ArchRProject (addGeneExpressionMatrix). Samples were derived from a mixture of both SOD1-G93A mice of a given stage and sex and control mice of the opposite sex. Each nucleus was assigned to the appropriate condition using counts of hSOD1 and sex-specific genes (*Xist*, *Uty*). Nuclei with ambiguous or conflicting marker gene expression were excluded.

### Multiome Sequencing: Clustering and Doublet Removal

Dimensionality reduction was performed on the ArchR project from above using the addIterativeLSI function on both the ATAC (clusterParams resolution = 0.2) and RNA modalities (clusterParams resolution = 0.2, useMatrix = “GeneExpressionMatrix”, depthCol = “Gex_nUMI”, varFeatures = 2500, firstSelection = “variable”, binarize = FALSE). A combined LSI representation was then computed from the ATAC and RNA LSI embeddings (addCombinedDims). UMAP embeddings were generated separately on the ATAC, RNA, and combined dimensions (addUMAP with minDist = 0.8), and clustering was performed on the combined LSI dimensions (addClusters with resolution = 0.4). Next, two clusters with high expression of marker genes that are known to be mutually exclusive were removed. The remaining clusters were renamed and then manually annotated in the cell_class metadata column using the expression of known marker genes.

### Multiome Sequencing: Cholinergic Neuron and Alpha Motor Neuron Subclustering

An ArchR project containing data from cholinergic neurons was created using the nuclei annotated as “Cholinergic Neurons” from the above analysis. Dimensionality reduction was performed on the resulting ArchR project using the addIterativeLSI function on both the ATAC (clusterParams resolution = 0.2) and RNA modalities (clusterParams resolution = 0.2, useMatrix = “GeneExpressionMatrix”, depthCol = “Gex_nUMI”, varFeatures = 2500, firstSelection = “variable”, binarize = FALSE). A combined LSI representation was then computed from the ATAC and RNA embeddings (addCombinedDims). UMAP embeddings were generated separately on the ATAC, RNA, and combined dimensions (addUMAP with minDist = 0.5), and clustering was performed on the combined LSI dimensions (addClusters with resolution = 0.5). One cluster with high expression of marker genes that are known to be mutually exclusive among cholinergic neurons was removed. The remaining clusters were renamed and then manually annotated in the cholinergic_type metadata column using the expression of known marker genes.

A similar approach was taken to the one above to create an ArchR project containing data from alpha motor neurons, with a few exceptions (addIterativeLSI: clusterParams resolution was left at the default value; addUMAP: minDist = 0.4). Clustering was performed using the ATAC-based LSI (addClusters with resolution = 1.2, nOutlier = 10), and the resulting clusters were manually annotated in the alpha_subtype metadata column using the expression of known marker genes and DAMN signature genes.

### Multiome Sequencing: Cholinergic Neuron Peak Calling with Downsampled Datasets

To generate a downsampled ArchR project of control cholinergic neurons, only nuclei annotated as “control” in the Stage metadata column were retained. A new metadata column was created by combining sample identity and cholinergic neuron type (e.g., Alpha MNs, Gamma MNs, Gamma* MNs, etc.). For each sample, the number of nuclei per cholinergic subtype was summarized, and the minimum number of nuclei among subtypes was determined. Nuclei were then randomly downsampled within each sample and subtype to the appropriate minimum count to ensure equal representation of cholinergic subtypes across samples. The resulting set of nuclei was used to subset the original cholinergic neuron ArchR project, producing a downsampled control dataset for downstream comparisons.

The downsampled control ArchR project was split into individual ArchR project by cholinergic neuron type (Alpha MNs, Gamma MNs, Gamma* MNs, Gad1+ cholinergic interneurons, Pitx2+ cholinergic interneurons, and Visceral MNs). For each subtype-specific project, pseudobulk replicates were generated by sample (addGroupCoverages with groupBy = “Sample”, minCells = 38, and minReplicates = 3). Peaks were then identified separately for each subtype using addReproduciblePeakSet with MACS2 (groupBy = “Sample”), and peak matrices were added with addPeakMatrix. These subtype-specific ArchR projects were subsequently used to quantify the number of peaks and to calculate the fraction of reads in peaks (FRIP) for each cholinergic neuron type.

The downsampled control ArchR project was used to create pseudobulk replicates (addGroupCoverages with groupBy = “cholinergic_type”, minCells = 38, and minReplicates = 3). Peaks were then called using addReproduciblePeakSet (groupBy = “cholinergic_type”), which used MACS2, and the peak matrix was added (addPeakMatrix). To quantify chromatin accessibility at single-nucleus resolution, the peak-by-cell accessibility matrix (PeakMatrix) was retrieved from the downsampled control ArchR project (getMatrixFromProject). For each nucleus, the number of accessible peaks was defined as the number of peaks with ≥1 fragment. To identify marker peaks across cholinergic neuron subtypes, differentially accessible peaks (marker peaks) were identified (getMarkerFeatures with groupBy = “cholinergic_type”, useMatrix = “PeakMatrix”, and testMethod = “wilcoxon”).

To compare chromatin accessibility changes with disease for motor neuron subtypes, alpha, gamma, gamma*, and visceral motor neurons were subset from the full cholinergic neuron ArchR project. Gamma and gamma* neurons were grouped as Pan-Gamma MNs, and individual ArchR projects were created for the three final subtypes: Alpha MNs, Pan-Gamma MNs, and Visceral MNs. For each sample/disease stage, the minimum number of nuclei among the three subtypes was computed, and nuclei were randomly downsampled within each sample/stage to match the appropriate minimum value, ensuring balanced representation across subtypes. Pseudobulk replicates were then created for each downsampled, subtype-specific ArchR project (addGroupCoverages with groupBy = “Stage” and minCells = 89). Peaks were called using addReproduciblePeakSet (groupBy = “Stage”), and the peak matrix was added (addPeakMatrix). Differentially accessible peaks were identified by comparing mid/end-stage SOD1-G93A motor neurons to controls for each subtype (getMarkerFeatures with groupBy = “Stage”, useGroups = “mid-late”, bgdGroups = “control”, useMatrix = “PeakMatrix”, testMethod = “wilcoxon”, and maxCells = 574).

### Multiome Sequencing: Alpha Motor Neuron Differentially Accessible Peaks with Disease

With the non-downsampled alpha motor neuron ArchR project from above, pseudobulk replicates were made (addGroupCoverages with groupBy = “Stage”), peaks were called (addReproduciblePeakSet with groupBy = “Stage”), and the peak matrices were added (addPeakMartrix). Differentially accessible peaks were identified by comparing mid/end-stage SOD1-G93A alpha motor neurons to controls (getMarkerFeatures with groupBy = “Stage”, useGroups = “mid-late”, bgdGroups = “control”, useMatrix = “PeakMatrix”, testMethod = “wilcoxon”, and maxCells = 1052).

### Multiome Sequencing: Identification of Positive Transcription Factor Regulators

Positive transcription factor (TF) regulators were identified for control skeletal motor neuron subtypes (alpha and pan-gamma), control alpha motor neuron subtypes (fast-firing, intermediate, and slow-firing), and alpha motor neuron disease status (non-DAMN, early DAMN, late DAMN). For each of these three analyses, an ArchR project containing the appropriate nuclei was created. With each ArchR project, pseudobulk replicates were made (addGroupCoverages with groupBy set appropriately), peaks were called (addReproduciblePeakSet with groupBy set appropriately), and the peak matrices were added (addPeakMartrix). Motif annotations were then added for each peak (addMotifAnnotations with motifSet = “cisbp”). Background peaks were generated (addBgdPeaks), and motif deviations and deviation z-scores were computed (addDeviationsMatrix with peakAnnotation = “Motif”). These data were aggregated by groups of interest (getGroupSE with useMatrix = “MotifMatrix” and groupBy set appropriately) and subsetted to just the deviation z-score data. For each motif, the maximum delta in z-score across groups (“maxDelta”) and the correlation between the motif accessibility and gene expression of the associated TF (correlateMatrices with useMatrix1 = “GeneExpressionMatrix”, useMatrix2 = “MotifMatrix”, and reducedDims = “LSI_Combined”) were calculated. The maxDelta information from above was added to the resulting correlation data frames. Motifs/TFs were labeled as “positive TF regulators” if they met the following criteria: correlation > 0.5, adjusted p-value < 0.01, and maxDelta above the 75th percentile.

### snRNA-seq/Multiome Sequencing: Label Transfer from Multiome to snRNA-seq Alpha Motor Neurons

The filtered count matrices containing the RNA data from the multiome experiments were imported (import10xFeatureMatrix function from ArchR). The resulting matrix was filtered to contain data from nuclei that are present in the alpha motor neuron ArchR project. The filtered matrix was used to create a Seurat object (CreateSeuratObject), and metadata (including alpha_subtype, which contains the firing properties/DAMN status information) from the alpha motor neuron ArchR project was added to the Seurat object (AddMetaData). The Seurat object was log-normalized (NormalizeData), and variable features were identified (FindVariableFeatures with selection.method = “vst” and features = 2000). To transfer the alpha_subtype labels to the RNA only alpha motor neuron Seurat object, transfer anchors were identified (FindTransferAnchors with reference set to the multiome RNA Seurat object, query set to the RNA only alpha motor neuron object, reference.assay = “RNA”, query.assay = “RNA”, dims = 1:50), and predictions were made (TransferData using the above anchorset, refdata set to the alpha_subtype metadata field from the multiome RNA Seurat object, dims = 1:50). These predictions were added as metadata (AddMetaData) to the RNA only alpha motor neuron Seurat object).

### snRNA-seq/Multiome Sequencing: In Silico Transcription Factor Perturbation with CellOracle

Transcription factor (TF) regulators of alpha motor neuron subtypes were identified using Python (v3.10) and CellOracle (v0.18.0)^50^, with analyses conducted separately for (1) control alpha motor neuron subtypes (fast-firing, intermediate, slow-firing) and (2) alpha motor neurons grouped by disease status (non-DAMN, early DAMN, late DAMN). To generate input for CellOracle, the snRNA-seq dataset (with subtype labels transferred from the multiome reference) was converted to AnnData format and pre-processed using Scanpy (v1.10.0). Pre-processing included gene filtering (minimum 1 count per gene using scanpy.pp.filter_genes), total count normalization (scanpy.pp.normalize_total), log transformation (scanpy.pp.log1p), selection of 2,000 highly variable genes (scanpy.pp.highly_variable_genes with flavor = “seurat”), PCA and neighbor graph construction (scanpy.tl.pca, scanpy.pp.neighbors), and UMAP embedding (scanpy.tl.umap). Discrete clusters that were not connected to the main alpha motor neuron population were removed.

To estimate pseudotime trajectories, a Pseudotime_calculator object (pt) was created for each analysis using CellOracle, with inputs including the UMAP coordinates and cluster labels. Lineages were manually defined (pt.set_lineage), and root cells were selected (pt.set_root_cells). Diffusion maps were computed with Scanpy (scanpy.tl.diffmap), and pseudotime values were calculated with get_pseudotime_per_each_lineage.

To prepare the data for regulatory network inference, a prebuilt mouse scATAC-seq-derived base GRN was imported (load_mouse_scATAC_atlas_base_GRN) as base_GRN, and Oracle objects were then initialized using raw count matrices, annotated cluster labels, and UMAP coordinates. The base GRN was then incorporated via oracle.import_TF_data. To reduce dimensionality and enable efficient k-nearest neighbors (KNN) imputation, PCA was performed (oracle.perform_PCA), and the number of components was selected based on a variance threshold, capped at 50. KNN imputation (oracle.knn_imputation) was applied using 2.5% of total cells as k (balanced = True, b_sight = k × 8, and b_maxl = k × 4).

Next, gene regulatory networks (GRNs) were inferred for each cell group (e.g., each control subtype or disease-status group) using oracle.get_links to generate Links objects (links). The resulting networks were filtered to retain high-confidence regulatory interactions (links.filter_links with p = 0.001, threshold_number = 10,000, weight = “coef_abs”). Group-specific TF dictionaries were then derived (oracle.get_cluster_specific_TFdict_from_Links) and used to fit the GRNs for downstream perturbation simulations (oracle.fit_GRN_for_simulation, alpha = 10, use_cluster_specific_TFdict = True).

To define a reference cell state trajectory, previously computed pseudotime values were transferred to the Oracle object (oracle.adata.obs[“Pseudotime”] = pt.adata.obs.Pseudotime). A Gradient_calculator object (gradient) was initialized using this pseudotime key. Cell density was estimated across a regular grid (gradient.calculate_p_mass with smooth = 0.8, n_grid = 40, and n_neighbors = 200), and low-density regions were filtered (gradient.calculate_mass_filter, min_mass = 6.3 or 8.3 depending on the analysis). The gene expression and pseudotime values were mapped onto the grid (gradient.transfer_data_into_grid with args = {“method”: “polynomial”, “n_poly”:3} for the control subtype analysis and oracle.get_cluster_specific_TFdict_from_Links for the disease analysis), and the gradient was computed (gradient.calculate_gradient), producing a reference vector field of unperturbed cell state transitions.

To evaluate the effect of TF perturbations, *in silico* knockout (KO) simulations were performed for all active regulatory TFs identified from the inferred GRNs (oracle.active_regulatory_genes). For each TF, CellOracle was used to simulate KO by setting the gene’s expression to zero (oracle.simulate_shift with n_propagation = 3). Cell state transitions were modeled using estimated transition probabilities (oracle.estimate_transition_prob with n_neighbors = 200, knn_random = True, and sampled_fraction = 1) and embedding shifts (oracle.calculate_embedding_shift with sigma_corr = 0.05). An Oracle_development_module object (dev) was instantiated for each TF, and the reference pseudotime gradient was loaded (dev.load_differentiation_reference_data). The simulated perturbation results were then imported (dev.load_perturb_simulation_data). Alignment between the KO vector field and the reference gradient was quantified by computing an inner product score (dev.calculate_inner_product), which was then discretized into 10 bins (dev.calculate_digitized_ip) to generate a perturbation score. Perturbation scores were analyzed using Oracle_systematic_analysis_helper, with statistical testing performed via calculate_positive_ps_p_value and calculate_negative_ps_p_value.

### MERFISH: Sample Collection and Imaging

For collection of fresh frozen spinal cord samples, mice were deeply anesthetized and then perfused with PBS. Following perfusion, the spinal cords were hydraulically extruded as described above or manually dissected. The tissue from each cord was cut into three or four small pieces centered around the cervical enlargement and another three to four small pieces centered around the lumbar enlargement. The cervical and lumbar tissue pieces were placed into respective plastic cryomolds that contained pre-chilled Optimal Cutting Temperature (OCT) compound. Using forceps, each cryomold was transferred into an isopentane and liquid nitrogen bath such that the cryomold made contact with the isopentane but was not submerged. The cryomolds maintained contact with the isopentane until the OCT had solidified. The samples were then stored at −80°C until the MERFISH experiments were performed.

The MERFISH experiments were conducted through the Vizgen MERSCOPE technology laboratory service. One MERSCOPE run contained only control, lumbar tissue (∼3-4 tissue sections from tissue pieces embedded together as described above), and the 18 additional runs contained both SOD1-G93A tissue at a given disease stage and age-matched control tissue (∼3-4 tissue sections each) per MERSCOPE run. Of these 18 runs, six contained early-stage SOD1-G93A tissue, six contained mid-stage SOD1-G93A tissue, and six contained end-stage SOD1-G93A tissue. Of the six runs per disease stage, two runs contained tissue centered around the cervical enlargement and four runs contained tissue centered around the lumbar enlargement. The full sample preparation user guide is available at https://vizgen.com/resources/fresh-and-fixed-frozen-tissue-sample-preparation/. For imaging, a custom gene panel for 140 genes was used. Additionally, *Apoe* and *Gfap* were included as sequential genes for all runs. For runs containing early or mid-stage SOD1-G93A tissue, *hSOD1-G93A* was also included as a sequential gene. The full instrument user guide is available at https://vizgen.com/resources/merscope-instrument/.

### MERFISH: Generation of a Custom Motor Neuron Segmentation Model

A custom motor neuron segmentation model was generated using the Cellpose3 (v3.1.0) graphical user interface.^109,110^ First, to generate training data, input images were created for each MERFISH experiment by plotting relevant transcripts as 10-pixel-diameter circles at their appropriate spatial coordinates with a custom Python script. One image included motor neuron/DAMN markers (*Chat*, *Prph*, *Slc5a7*, *Atf3*) in green, and a second image included non-motor neuron markers (*Aldh1l1*, *Aqp4*, *Cx3cr1*, *Gad1*, *Mog*, *Slc17a6*, *Slc6a5*, *Trem2*) in red. These input images were used with the Vizgen Post-processing Tool (vpt, v1.3.0) to extract 600 × 600 µm image patches from selected locations across spinal cord tissue sections and experiments (vpt extract-image-patch). The resulting image patches were used to manually trace motor neurons using the Graphic app (Picta) on an iPad (Apple). The manually traced motor neuron masks were post-processed using a custom ImageJ macro to create Cellpose-compatible segmentation masks. The image patches and masks were then used to train a new Cellpose model (initial model: cyto3, chan to segment: green, chan2: red, learning_rate: 0.1, weight_decay: 0.0001, n_epochs: 500). The resulting model was validated visually on held-out images to confirm accurate segmentation of large cholinergic motor neurons and was used for downstream segmentation tasks.

### MERFISH: Cell Segmentation, Transcript Partitioning, and Cell Metadata Calculation

Cell segmentation, transcript partitioning, and cell metadata calculation were performed using the Vizgen Post-processing Tool (vpt, v1.3.0) with the Cellpose2 plugin (vpt-plugin-cellpose2, v1.0.1).^111^ To segment motor neurons and other cell types, a custom two-task segmentation algorithm was used. For motor neuron segmentation, the custom model described above was applied to full-experiment input images showing motor neuron and non-motor neuron transcript locations. For segmentation of other cells, a standard Cellpose2 “nuclei” model was applied using the DAPI channel. Segmentation outputs from the two tasks were combined using a “harmonize” fusion strategy to produce a complete, non-overlapping set of cell boundaries. This segmentation algorithm was used to identify cell boundaries for each MERFISH experiment (vpt run-segmentation with a tile size of 7000 and tile overlap of 900). After segmentation, transcripts were partitioned into cells (vpt partition-transcripts), and then cell metadata was calculated (vpt derive-entity-metadata, vpt sum-signals).

### MERFISH: Data Pre-Processing and Initial Clustering

Subsequent MERFISH analyses were performed in Python (v3.9.21) using the Scanpy package (v1.10.3).^112^ For each MERFISH experiment, an AnnData object was created by combining gene expression data, cell metadata, and signal intensities for the sequential genes (*Apoe*, *Gfap*, and *hSOD1-G93A*). To account for multiple tissue sections within a single experiment, each cell was assigned to a specific section based on its spatial coordinates. Cells that did not fall within any defined tissue section were excluded. For cells within valid sections, additional metadata was added, including section ID, slide ID, anatomical region, disease stage, and experiment batch (VS119 or VS223). For each MERFISH experiment, genes labeled as “blank” were removed as well as cells with 20 or fewer transcripts or five or fewer detected genes. The filtered AnnData objects were then concatenated into a single dataset with raw counts stored in .X and duplicated in .layers[“counts”]. Additionally, a few low-quality tissue sections were removed based on manual inspection.

To account for cell size differences, sequential gene signals and raw gene expression values were normalized by cell volume. Total gene expression for each cell was then normalized to 250 (scanpy.pp.normalize_total), and expression values were log-transformed with a pseudocount (scanpy.pp.log1p). The log-normalized gene expression values were subsequently converted to z-scores (scanpy.pp.scale with max_value = 10), and principal component analysis (PCA) was performed for dimensionality reduction (scanpy.tl.pca). A batch-balanced k-nearest neighbors graph was then constructed using BBKNN (scanpy.external.pp.bbknn), with experiment batch (VS119 or VS223) specified as the batch key. This graph was used to compute a UMAP embedding for visualization (scanpy.tl.umap). Leiden clustering was then performed (scanpy.tl.leiden with resolution = 0.5, flavor = “igraph”, and n_iterations = 2), and the resulting clusters were manually annotated based on the expression of cell class marker genes and their spatial locations.

### MERFISH: Cholinergic Neuron and Alpha Motor Neuron Subclustering

To subcluster cholinergic neurons, cells labeled as “Cholinergic Neurons” were subset from the full AnnData object. Batch correction across experiment sets (VS119 and VS223) was performed using BBKNN (scanpy.external.pp.bbknn), followed by UMAP embedding (scanpy.tl.umap with min_dist = 0.1) and Leiden clustering (scanpy.tl.leiden with resolution = 1.0, flavor = “igraph”, and n_iterations = 2). Clusters enriched for *Mog* expression were identified and evaluated as putative peri-motor neuron oligodendrocytes. To assess this, the volume distribution of these cells was compared to that of high-confidence oligodendrocytes from the full dataset using violin plots. Interquartile range (IQR)-based whisker bounds were used to define the expected volume range for oligodendrocytes. Cells falling within this range were reclassified as oligodendrocytes, while those outside the range were removed. The remaining cholinergic neurons were then reanalyzed: BBKNN and UMAP were recomputed, and Leiden clustering was repeated with increased resolution (resolution = 1.2). The resulting clusters were manually annotated using the expression of known cholinergic neuron marker genes. Finally, cells with a volume less than 1000 were removed to exclude low-volume cholinergic neurons. To subcluster alpha motor neurons, cells labeled as “Alpha MNs” were subset from the cholinergic neuron dataset. BBKNN (scanpy.external.pp.bbknn), UMAP (scanpy.tl.umap), and Leiden clustering (scanpy.tl.leiden with resolution = 0.45, flavor = “igraph”, and n_iterations = 2) were then applied to identify subclusters.

### MERFISH: Label Transfer from snRNA-seq to MERFISH Alpha Motor Neurons

To annotate alpha motor neuron subtypes in the MERFISH dataset, label transfer was performed using the alpha motor neuron snRNA-seq dataset in AnnData format as a reference. Subtype labels (“Fast-Firing”, “Intermediate”, “Slow-Firing”, “Early DAMN”, and “Late DAMN”) were originally defined in the multiome dataset and subsequently transferred to the snRNA-seq dataset, as described above, prior to use as the reference. Both the MERFISH and snRNA-seq datasets were filtered to include only shared genes, and a batch label was added to distinguish their origins. The datasets were concatenated and jointly embedded using scVI (scvi.model.SCVI, scvi-tools v1.1.6.post2).^113,114^ The SCVI model was trained on the combined dataset with batch correction (SCVI.setup_anndata and SCVI.train). Cell type label transfer was then performed using SCANVI, a semi-supervised extension of scVI.^114,115^ The SCANVI model was initialized from the pretrained SCVI model (SCANVI.from_scvi_model) and trained using the snRNA-seq annotations as the labeled reference and the MERFISH cells as unlabeled (SCANVI.train with max_epochs = 400). Predicted cell type labels were inferred for each MERFISH alpha motor neuron (SCANVI.predict) and stored in the alpha motor neuron AnnData object.

### MERFISH: Alpha Motor Neuron Differential Expression Analysis with Disease

To identify differentially expressed genes in alpha motor neurons, a set of disease-associated genes previously defined in the snRNA-seq analysis (control vs. end-stage; adjusted p-value < 0.01) was intersected with the MERFISH gene panel. The resulting gene set was used to subset the MERFISH alpha motor neuron dataset, and raw expression values were retrieved from the counts layer. Expression was then normalized by cell volume, scaled to 250 total counts per cell (scanpy.pp.normalize_total), and log-transformed with a pseudocount (scanpy.pp.log1p). Pairwise differential expression analysis was performed between control vs. mid-stage and control vs. end-stage alpha motor neurons using two-sided Mann-Whitney U tests. Fold changes were computed on the linear (non-log-transformed) scale and then log2-transformed, and adjusted p-values were calculated using the Benjamini-Hochberg procedure (statsmodels.stats.multitest.multipletests).

### MERFISH: Dataset Refinement and Spatial Standardization

An updated version of the full MERFISH dataset was generated to refine and extend cell type annotations. To do so, subsets of the original AnnData object were selectively replaced: peri-motor neuron oligodendrocytes were reassigned to the “Oligodendrocytes” class, and all cholinergic neurons were removed and replaced with the filtered dataset excluding low-volume cells and oligodendrocyte contaminants. Alpha motor neurons were replaced with a label-transferred dataset containing predicted subtypes, and a DAMN_status column was added to classify them as “DAMN” or “Non-DAMN.” To standardize section orientation, as done in Sun et al.^116^, spatial coordinates were mean-centered and rotated using section-specific angles determined by visual inspection. Updated coordinates were stored in adata.obsm[“spatial”].

### MERFISH: Tissue Section Filtering for Spatial Analyses

For analyses sensitive to tissue integrity, such as spatial neighborhood and relative abundance analyses, sections were further filtered to exclude sections with tears. Each tissue section was manually reviewed and labeled as “keep,” “remove,” “left,” or “right.” Sections labeled “keep” were retained in full, while left and right halves were extracted from “left” and “right” sections, respectively. Cells from “keep,” left-filtered, and right-filtered sections were merged to generate a high-quality spatial dataset for downstream analysis.

### MERFISH: Reactive Glial Cell Classification

To identify reactive glial cells across disease stages, per-experiment expression thresholds were computed based on control-stage tissue. For each slide (excluding one control-only slide with no disease tissue), the 95th percentile of *Apoe* expression among control microglia/macrophages and of *Gfap* expression among control astrocytes was calculated. These values were computed using high-pass filtered and normalized expression values. Cells were labeled based on these control-derived thresholds: microglia/macrophages with elevated *Apoe* were classified as reactive, and astrocytes with elevated *Gfap* were classified as reactive or white matter astrocytes.

### snRNA-seq: Cell-Cell Communication Analysis with CellChat

To identify disease-associated ligand-receptor interactions, we performed cell-cell communication analysis using the CellChat package (v2.2.0)^77^ in R (v4.4.2). An expression matrix and metadata were first exported from a final snRNA-seq Seurat object that contained all high-quality data and annotations. Specifically, the normalized expression data were obtained with the Seurat function GetAssayData (assay = “RNA”, slot = “data”), and the full metadata were extracted from the meta.data. These were saved as .rds files and used as input for CellChat. For the CellChat analysis, the dataset was restricted to three cell types of interest (alpha motor neurons, astrocytes, and microglia/macrophages) and two conditions (control and end-stage).

For each condition (control and end-stage), a CellChat object was constructed from the filtered count matrix and cell metadata (createCellChat with group.by set to cell type labels). The ligand-receptor interaction database was restricted to the secreted signaling subset of CellChatDB.mouse (subsetDB with search = “Secreted Signaling” and key = “annotation”). For each CellChat object, the expression data were preprocessed (subsetData, identifyOverExpressedGenes with do.fast = FALSE, and identifyOverExpressedInteractions), and communication probabilities were computed (computeCommunProb with type = “truncatedMean” and trim = 0.1). Subsequently, signaling pathways were inferred (computeCommunProbPathway), the communication network was aggregated (aggregateNet), and network centrality scores were calculated (netAnalysis_computeCentrality).

The control and disease CellChat objects were then merged (mergeCellChat) for a comparative analysis. Differential expression between conditions was assessed using identifyOverExpressedGenes with the disease condition set as the positive dataset (only.pos = FALSE, thresh.pc = 0.1, thresh.fc = 0.1, thresh.p = 0.05, group.DE.combined = FALSE, do.fast = FALSE. The differentially expressed gene information was mapped onto the inferred cell-cell communications (netMappingDEG), and ligand-receptor pairs showing significantly increased or decreased signaling in disease relative to control were extracted (subsetCommunication; either datasets set to the positive, disease dataset, ligand.logFC = 0.1, receptor.logFC = NULL or datasets set to the negative, control dataset, ligand.logFC = −0.1, receptor.logFC = NULL). To avoid ambiguity, ligand-receptor interactions appearing in both up- and downregulated sets were removed. Visualization of differentially regulated interactions was performed using the netVisual_bubble function, and bubble plots were generated for each source population (alpha motor neurons, astrocytes, microglia/macrophages) against other potential target populations.

### Partitioned Heritability and H-MAGMA Analyses

Partitioned heritability was estimated with Linkage Disequilibrium (LD) Score Regression v1.0.1 via the command line to evaluate enrichment of genome-wide associated study (GWAS) signals within differentially accessible chromatin regions.^88,117^ Differentially accessible peaks in mid/end-stage SOD1-G93A vs. control alpha motor neurons (FDR < 0.05) were lifted over from mm10 to hg19 genome using the bnMapper function from the bx-python package v0.12.0.^118^ After conversion, 250 bp padding was added to both sides of each peak, and blacklisted hg19 regions were removed.^89,119^ Annotation files were generated by identifying all HapMap3 single nucleotide polymorphisms (SNPs)^120^ located within the lifted over peaks. LD scores were then calculated using default settings (LD window: 1cM, reference panel: 1000 Genomes European Phase 3.^121^ Partitioned heritability was estimated by including LD scores from both the baseline model and SNPs within all lifted-over differentially accessible peaks in the regression.^88^ GWAS summary statistics for ALS and other disorders were used to compute trait-specific partitioned heritability.^90–93^ Enrichment was defined as the proportion of heritability explained by SNPs in the annotation (i.e., lifted-over differentially accessible peaks) divided by the proportion of reference SNPs within the annotation. Enrichment p-values were FDR-corrected across the four GWAS datasets.

To identify putative genes underlying the ALS GWAS signal enrichment within lifted-over differentially accessible regions, H-MAGMA was performed using the most recent ALS GWAS, following the developer’s instructions.^94,95^ H-MAGMA extends conventional Multi-marker Analysis of GenoMic Annotation (MAGMA) by incorporating Hi-C-derived 3D chromatin interaction data to link intergenic SNPs to their potential target genes.^122^ Required input files, including exon and promoter coordinates, SNP coordinates, their overlaps, and adult brain Hi-C data, were downloaded from Zenodo (https://doi.org/10.5281/zenodo.5503876). Loop anchors in the Hi-C dataset were intersected with promoter regions to define anchor 1 sites. Intergenic SNPs located in the corresponding anchor 2 sites were then assigned to the gene whose promoter overlapped anchor 1. The resulting SNP-gene annotation table was then used for gene-level analysis with the SNP-wise model in MAGMA v1.10. The final H-MAGMA output, containing *Z* statistics for each gene, was intersected with genes in the lifted-over differentially accessible peaks to identify putative ALS risk genes within regions of altered chromatin accessibility.

## Data and code availability

Raw and processed sequencing data as well as original code will be made available upon publication.

---

**Figure S1:**
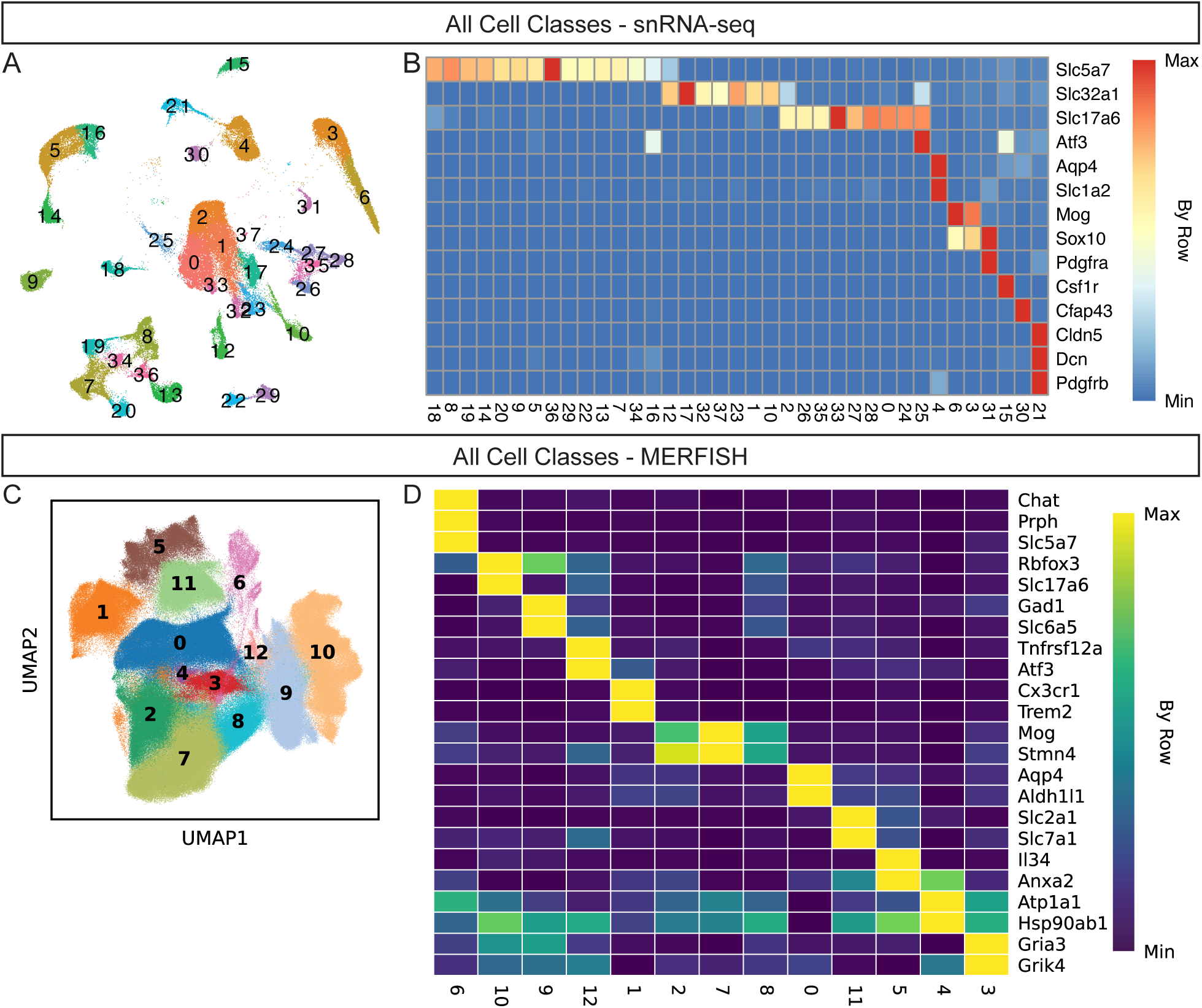
Identification of major cell classes by snRNA-seq and MERFISH. **(A)** UMAP representation of all snRNA-seq nuclei labeled by cluster. **(B)** Heatmap showing average expression levels of depicted genes for each cluster from (A) min/max-normalized by row. These data were used to assign cell class annotations for the snRNA-seq data. **(C)** UMAP representation of MERFISH cells labeled by cluster. **(D)** Heatmap showing average expression levels of depicted genes for each cluster from (C) min/max-normalized by row. These data were used to assign cell class annotations for the MERFISH data

**Figure S2:**
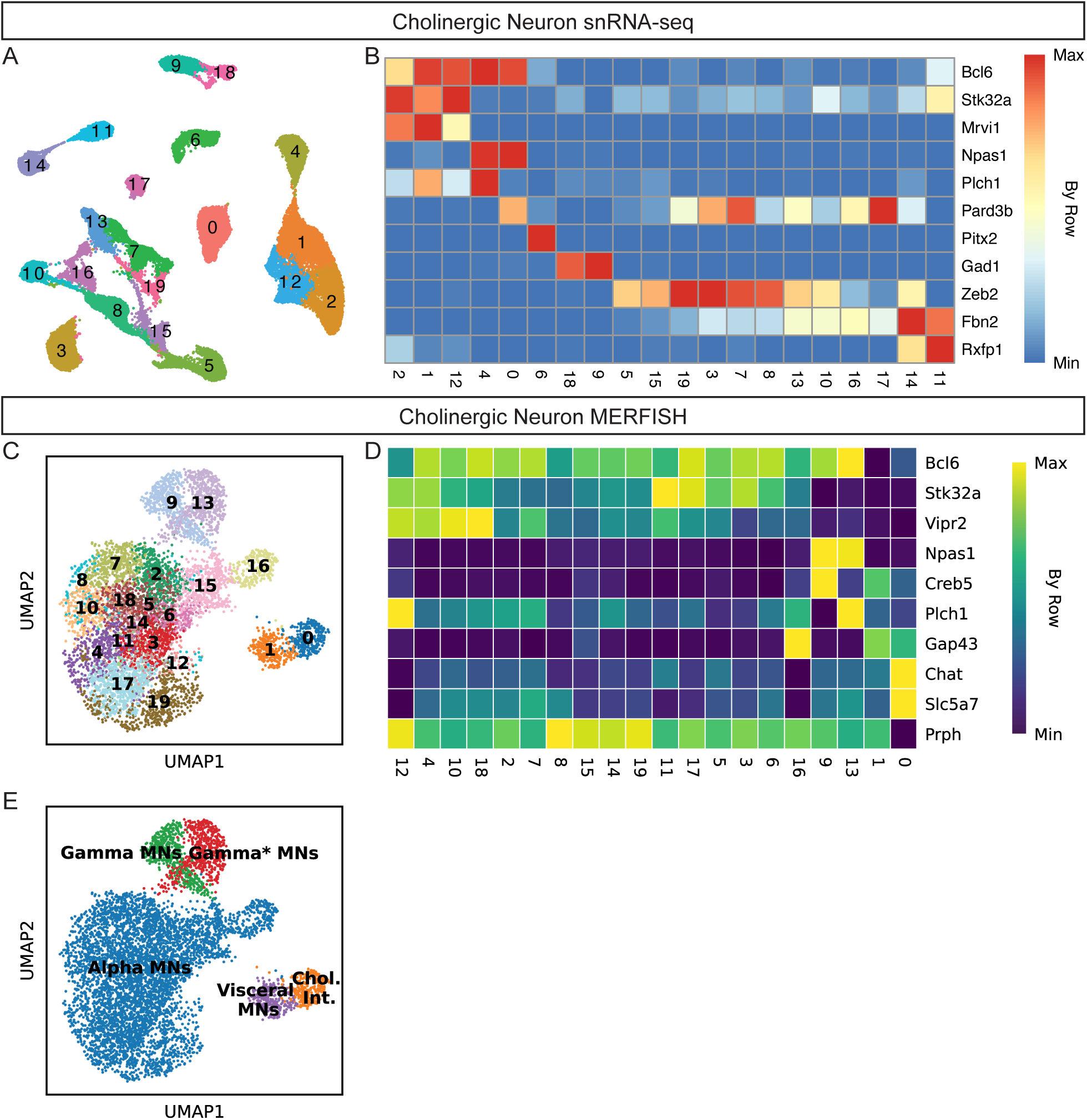
Identification of cholinergic neuron types by snRNA-seq and MERFISH. **(A)** UMAP representation of cholinergic neurons labeled by cluster. **(B)** Heatmap showing average expression levels of depicted genes for each cluster from (A) min/max-normalized by row. These data were used to assign cholinergic neuron type annotations for the snRNA-seq data. **(C)** UMAP representation of MERFISH cholinergic neurons labeled by cluster. **(D)** Heatmap showing average expression levels of depicted genes for each cluster from (C) min/max-normalized by row. These data were used to assign cholinergic neuron type annotations for the MERFISH data. **(E)** UMAP representation of the MERFISH cholinergic neurons from (C) labeled by cholinergic neuron type.

**Figure S3:**
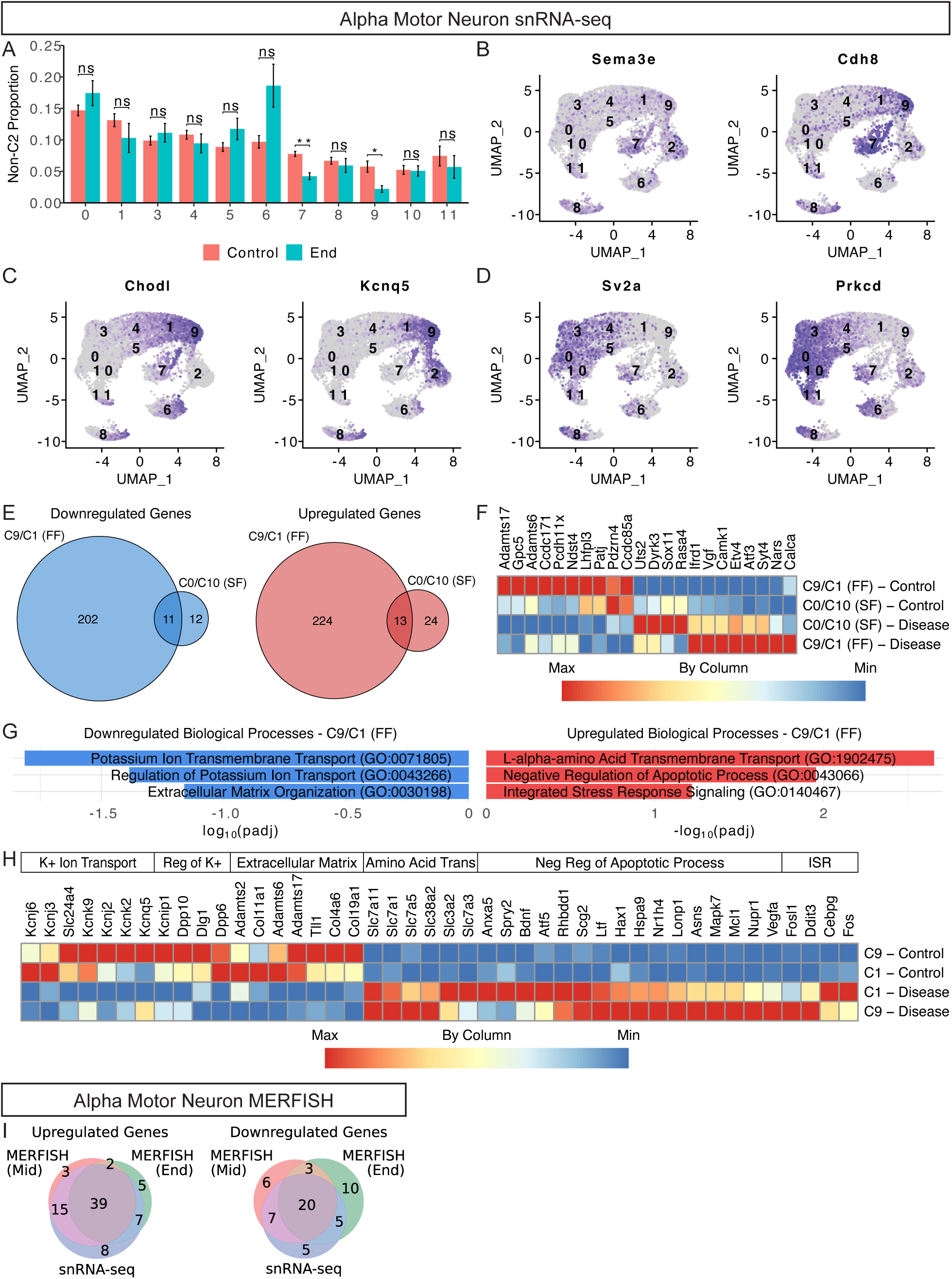
Subtype-specific responses of alpha motor neurons to disease. **(A)** Bar plots of non-cluster 2 alpha motor neuron proportions by cluster, comparing control and end-stage conditions. Bars show group means ± standard error across experiments (n = 11 control experiments, n = 6 end-stage experiments). Asterisks indicate significance (Welch’s t-test per cluster with Benjamini-Hochberg correction): n.s. = not significant; * padj ≤ 0.05; ** padj ≤ 0.01. **(B-D)** UMAP plots showing expression of representative marker genes of gluteus/shoulder-innervating **(B)**, fast-firing **(C)**, and slow-firing **(D)** alpha motor neurons. Each dot represents a single alpha motor neuron, colored by gene expression (darker purple color = higher expression). **(E)** Venn diagrams of genes significantly downregulated (left) or upregulated (right) in pre-DAMN fast-firing (clusters 9 and 1) and slow-firing (clusters 0 and 10) alpha motor neurons in disease (mid or end-stage) compared to control (Wald test with Benjamini-Hochberg correction; padj < 0.01). **(F)** Heatmap showing average expression levels of selected differentially expressed genes in both fast-firing and slow-firing pre-DAMN alpha motor neurons with disease. Data are min/max-normalized by column. **(G)** Bar plots showing log_10_(padj) or −log_10_(padj) for selected downregulated (left) and upregulated (right) GO biological processes with disease in pre-DAMN fast-firing alpha motor neurons. **(H)** Heatmap showing average expression levels of selected differentially expressed genes in pre-DAMN fast-firing alpha motor neurons (clusters 9 and 1) with disease. Data are min/max-normalized by column. **(I)** Venn diagrams showing significantly upregulated (top) and downregulated (bottom) genes in alpha motor neurons across MERFISH and snRNA-seq datasets. MERFISH results compare mid-or end-stage alpha motor neurons to controls (two-sided Mann-Whitney U test with Benjamini-Hochberg correction; FDR < 0.01). For snRNA-seq, only differentially expressed genes from Figure 1G/Table S1A that are present in the MERFISH panel were included.

**Figure S4:**
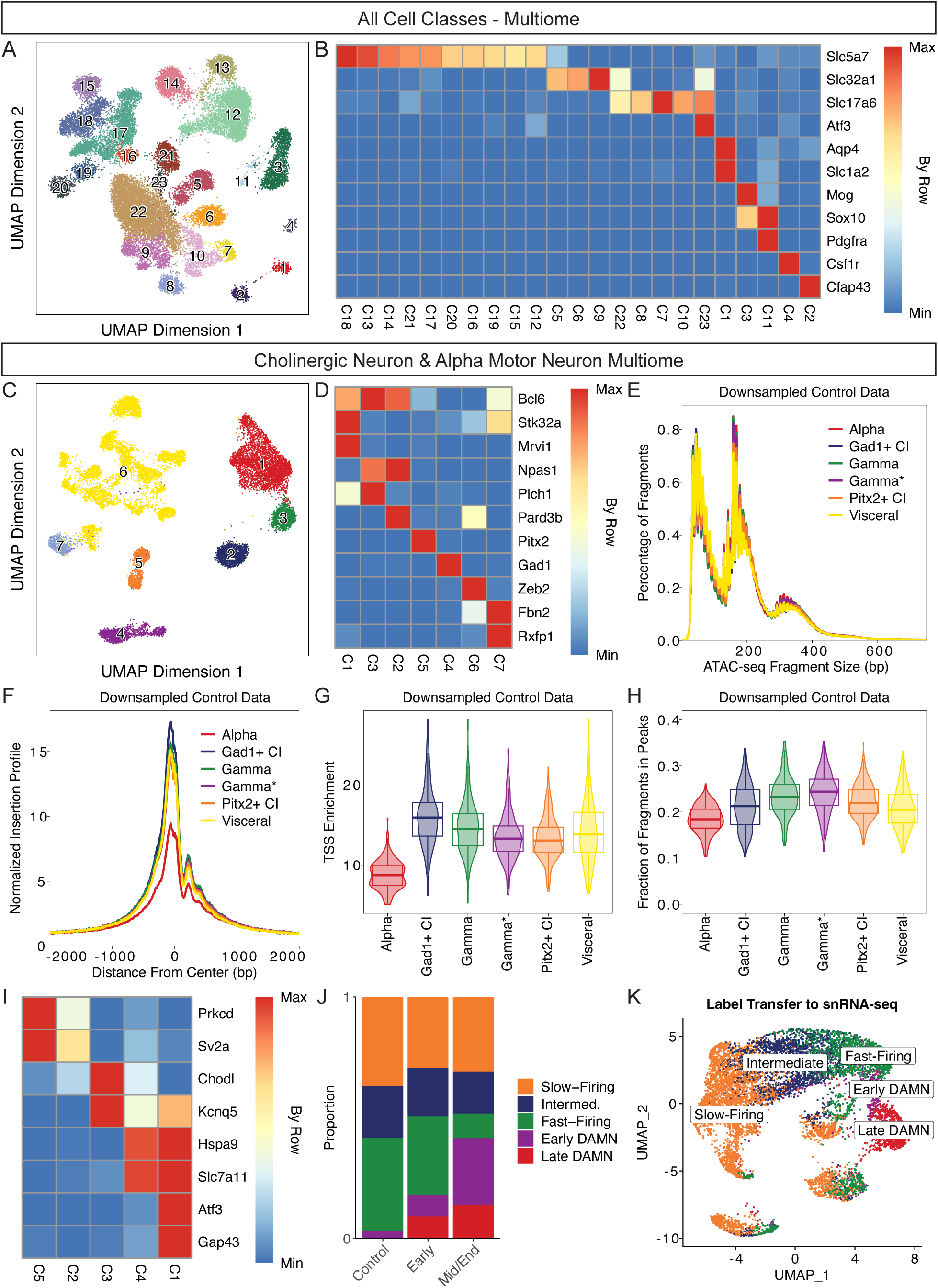
Identification of major cell classes and cholinergic neuron types by paired snATAC/snRNA-seq. **(A)** UMAP representation of all paired snATAC/snRNA-seq profiles labeled by cluster. **(B)** Heatmap showing average expression levels of depicted genes for each cluster from (A) min/max-normalized by row. These data were used to assign cell class annotations. **(C)** UMAP representation of cholinergic neuron paired snATAC/snRNA-seq profiles labeled by cluster. **(D)** Heatmap showing average expression levels of depicted genes for each cluster from (C) min/max-normalized by row. These data were used to assign cholinergic neuron type annotations. **(E)** Fragment size distributions for each cholinergic type using downsampled control snATAC-seq data. **(F)** TSS enrichment profiles for each cholinergic type using downsampled control snATAC-seq data. **(G)** Violin plots showing the distribution of TSS enrichment scores across cholinergic neuron types in downsampled control data with box plots overlaid. **(H)** Violin plots showing the distribution of Fraction of Reads/Fragments in Peaks (FRIP) scores across cholinergic neuron types in downsampled control data with box plots overlaid. FRIP was calculated separately for each cholinergic neuron type using the subtype-specific peak sets from Figure 3E. **(I)** Heatmap showing average expression levels of depicted genes for each alpha motor neuron cluster from Figure 3G min/max-normalized by row. These data were used to assign alpha motor neuron subtype annotations. **(J)** Stacked bar plots showing the proportion of nuclei assigned to each alpha motor neuron subtype in control and disease conditions. **(K)** UMAP representation of snRNA-seq alpha motor neurons labeled by alpha motor neuron subtype after label transfer from the multiome data.

**Figure S5:**
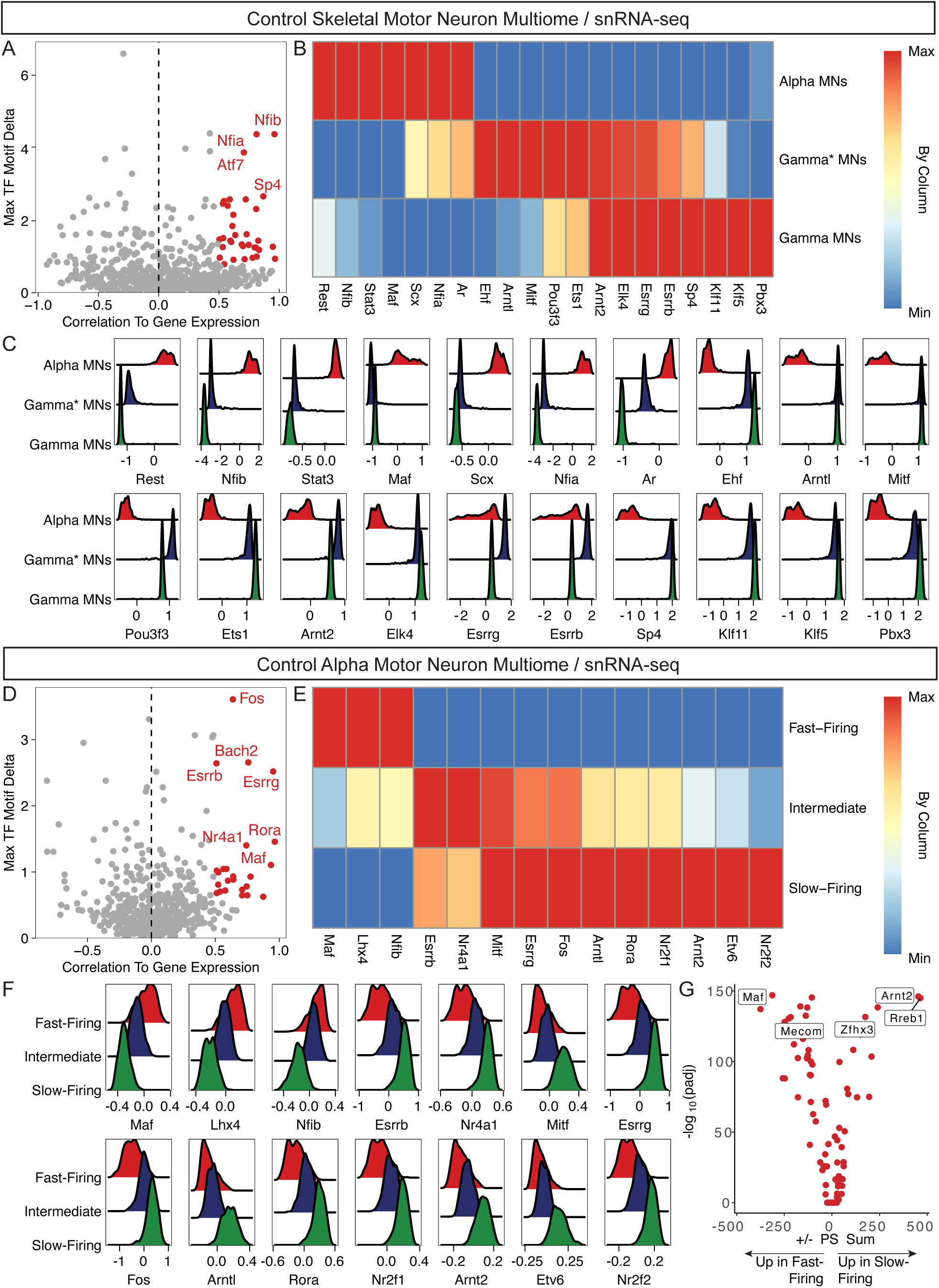
Transcription factor (TF) regulators of skeletal motor neuron subtypes. **(A)** Dot plot showing the identification of positive TF regulators of alpha, gamma*, and gamma motor neurons with ArchR. TFs with maximum differences in chromVAR deviation z-score in the top quartile of all TFs and a correlation of greater than 0.5 are plotted in red and are candidate positive TF regulators. **(B)** Heatmap showing average expression levels of selected differentially expressed TFs between alpha and pan-gamma motor neurons by snRNA-seq. Data are min/max-normalized by column. **(C)** Distributions of chromVAR deviation scores of TFs from (B) for alpha, gamma*, and gamma motor neurons. **(D)** Dot plot showing the identification of positive TF regulators of alpha motor neuron functional subtypes (fast-firing, intermediate, slow-firing) with ArchR. TFs with maximum differences in chromVAR deviation z-score in the top quartile of all TFs and a correlation of greater than 0.5 are plotted in red and are candidate positive TF regulators. **(E)** Heatmap showing average expression levels of selected differentially expressed TFs between fast-firing and slow-firing alpha motor neurons by snRNA-seq after label transfer from the multiome data. Data are min/max-normalized by column. **(F)** Distributions of chromVAR deviation scores of TFs from (E) for fast-firing, intermediate, and slow-firing alpha motor neurons. **(G)** CellOracle *in silico* knockout results of 56 TFs in fast-firing and slow-firing alpha motor neurons summarized as a volcano plot with the sum of positive or negative perturbation scores on the x-axis and −log_10_(padj) on the y-axis.

**Figure S6:**
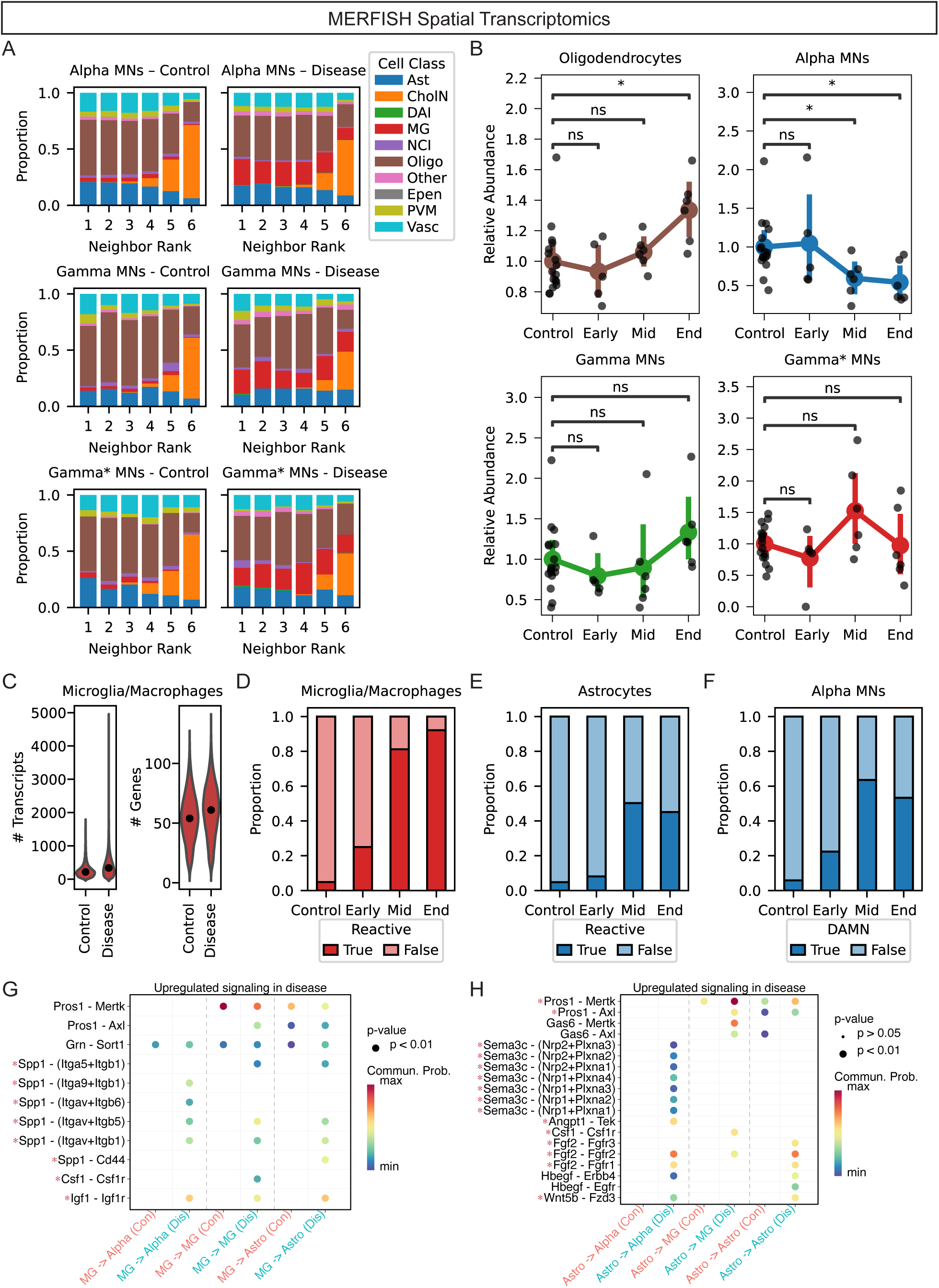
Extended spatial and compositional dynamics of spinal cord cell types with disease in the SOD1-G93A mouse. **(A)** Stacked bar plots showing the cell class composition of the six nearest neighbors to alpha (top), gamma (middle), and gamma* (bottom) motor neurons in control (left) and disease (right) conditions (Ast: Astrocytes, CholN: Cholinergic Neurons, DAI: Disease-Associated Interneurons, MG: Microglia/Macrophages, NCI: Non-Cholinergic Interneurons, Oligo: Oligodendrocytes, Epen: Putative Ependymal Cells, PVM: Putative Perivascular/Meningeal Cells, Vasc: Putative Vascular Cells). Microglia/macrophages are significantly enriched at all neighbor positions for all skeletal motor neuron subtypes in disease compared to control (Fisher’s Exact Test, Bonferroni-adjusted padj ≤ 0.01). **(B)** Strip and point plots showing the abundance of oligodendrocytes (top left), alpha (top right), gamma (bottom left), and gamma* (bottom right) skeletal motor neurons relative to interneurons across control and disease conditions. Each point represents the average value from tissue sections of a MERFISH experiment (n ≥ 5 experiments per condition). Values are scaled to the mean relative abundance in control samples. Colored points and lines indicate cell type means ± 95% confidence intervals. Asterisks indicate significance relative to control (one-sided Welch’s t-test, Bonferroni-adjusted): n.s. = not significant; * padj ≤ 0.05. **(C)** Violin plots showing the distribution of unique transcripts (left) and genes (right) detected per cell in microglia/macrophages from control and disease conditions. Each violin represents the distribution across individual cells. Black points indicate median values: 218 transcripts and 54 genes in control, and 338 transcripts and 61 genes in disease. **(D)** Stacked bar plots showing the proportion of reactive and non-reactive microglia/macrophages across control and disease conditions. **(E)** Stacked bar plots showing the proportion of reactive/white matter and other astrocytes across control and disease conditions. **(F)** Stacked bar plots showing the proportion of DAMN and non-DAMN alpha motor neurons across control and disease conditions. **(G-H)** Bubble plots of ligand-receptor interactions upregulated in disease with microglia/macrophages **(G)** and astrocytes **(H)** as the sender cell types. Bubble size indicates statistical significance, and color reflects communication probability. Red stars indicate ligand-receptor interactions where the ligand is significantly upregulated in microglia/macrophages or astrocytes, respectively, at end-stage by the snRNA-seq.

**Figure S7:**
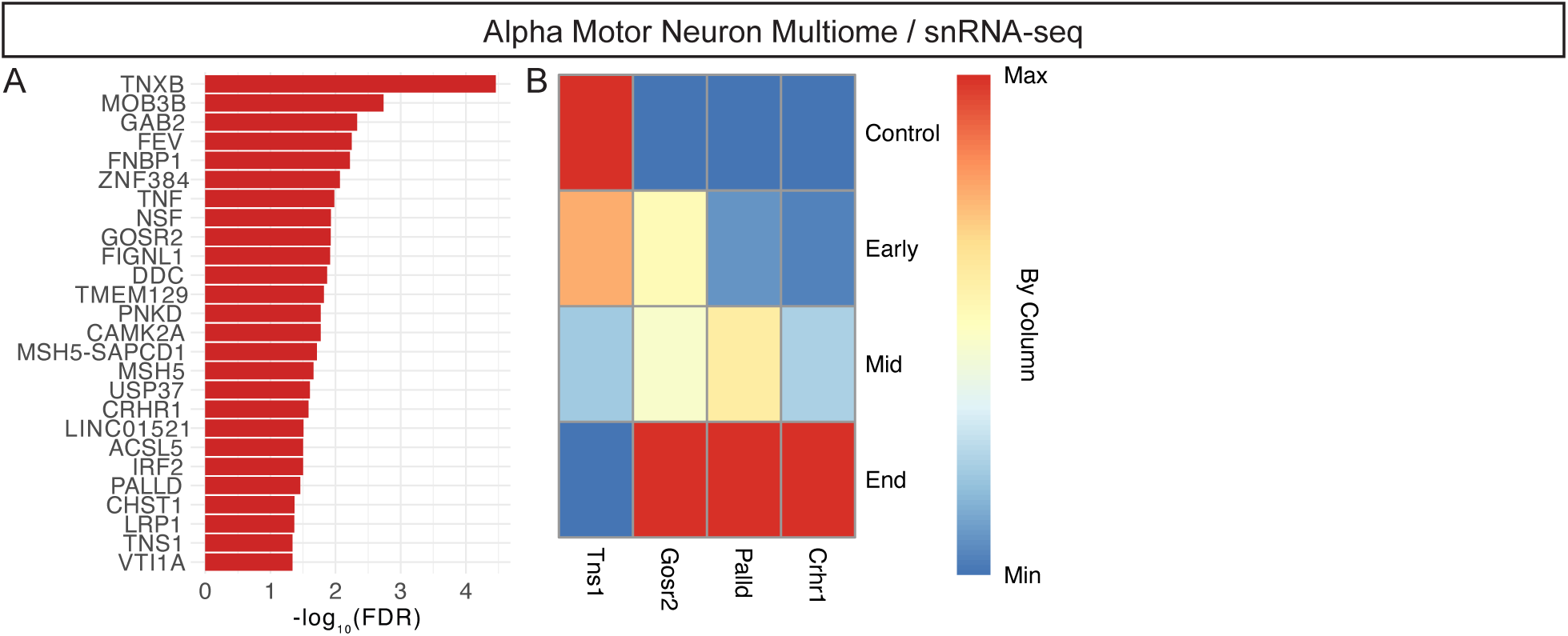
Candidate ALS-associated genes from H-MAGMA. **(A)** Bar plot showing candidate ALS-associated genes based on H-MAGMA analysis and alpha motor neuron differential chromatin accessibility with disease. Bars represent −log_10_(FDR) values for each gene (FDR < 0.05). Gene-level associations were derived by mapping non-coding SNPs to target genes using chromatin interaction profiles. **(B)** Heatmap showing average expression levels of differentially expressed genes from (A) throughout disease progression in alpha motor neurons by snRNA-seq. Data are min/max-normalized by column.

## Notes

https://spinalcordatlas.org

